# Structure of the Ion Channel Kir7.1 and Implications for its Function in Normal and Pathophysiologic States

**DOI:** 10.1101/2024.06.07.597981

**Authors:** Alys Peisley, Ciria C. Hernandez, Naima S. Dahir, Laura Koepping, Ashleigh Raczkowski, Min Su, Masoud Ghamari-Langroudi, Xinrui Ji, Luis E. Gimenez, Roger D. Cone

## Abstract

Hereditary defects in the function of the Kir7.1 in the retinal pigment epithelium are associated with the ocular diseases retinitis pigmentosa, Leber congenital amaurosis, and snowflake vitreal degeneration. Studies also suggest that Kir7.1 may be regulated by a GPCR, the melanocortin-4 receptor, in certain hypothalamic neurons. We present the first structures of human Kir7.1 and describe the conformational bias displayed by two pathogenic mutations, R162Q and E276A, to provide an explanation for the basis of disease and illuminate the gating pathway. We also demonstrate the structural basis for the blockade of the channel by a small molecule ML418 and demonstrate that channel blockade in vivo activates MC4R neurons in the paraventricular nucleus of the hypothalamus (PVH), inhibiting food intake and inducing weight loss. Preliminary purification, and structural and pharmacological characterization of an in tandem construct of MC4R and Kir7.1 suggests that the fusion protein forms a homotetrameric channel that retains regulation by liganded MC4R molecules.

## INTRODUCTION

Inward-rectifier K^+^ channels play a vital role in cell function by allowing K^+^ ions to enter cells, particularly at hyperpolarized membrane potentials, thus establishing and preserving the resting membrane potential [1, 2]. Among these channels, Kir7.1 is distinguished by its expression in various epithelial tissues, notably in the retinal pigment epithelium (RPE), crucial for maintaining the blood-retinal barrier [3-5]. Mutations in Kir7.1 are associated with several ocular disorders, including retinitis pigmentosa, Leber congenital amaurosis, and snowflake vitreal degeneration, with the severity of visual impairment correlating with the extent of channel dysfunction [6, 7]. Kir7.1 also exhibits a unique feature reminiscent of the regulation of inward rectifier Kir6.2 by the multi-membrane spanning sulfonylurea receptor [8]. Data from studies in cell culture and hypothalamic slice preparations suggest that Kir7.1 can be directly regulated by a GPCR in a G-protein-independent manner [9], playing a crucial role in the rapid depolarization of melanocortin 4 receptor (MC4R) neurons in response to its ligand, α-melanocyte stimulating hormone (α-MSH), and hyperpolarization in response to its ligand agouti related protein (AgRP). Thus Kir7.1, the most divergent member of the family of inward rectifier channels, displays a variety of unusual functions in diverse cell types.

K^+^ channels exhibit three distinct states: resting, activated, and inactivated [10, 11]. Typically, they remain closed in the resting state, open upon stimulation, and return to a non-conductive state afterward. Using cryo-electron microscopy, we analyze here the structures of full-length Kir7.1 under various conditions, including wild-type apo, bound to phosphatidylinositol 4,5-bisphosphate (PIP_2_) or treated with the channel blocker ML418 [12], as well as the mutated forms R162Q and E276A associated with retinitis pigmentosa [7]. Our structural data confirmed that the channel blocker ML418 binds to Kir7.1. We then studied its role in regulating MC4R neurons, feeding behavior, and body weight. In addition, we purified and characterized MC4R-Kir7.1 fusion proteins to understand the structural characteristics that govern the interaction between MC4R and Kir7.1.

## RESULTS

### Protein expression and purification of human Kir7.1 and R162Q and E276A mutants

The human wild-type Kir7.1 channel and its mutant counterparts, R162Q and E276A, were produced in Sf9 insect cells infected with recombinant baculovirus harboring GB1-tagged Kir7.1 constructs. The size exclusion chromatography (SEC) profile of affinity-purified Kir7.1 (Figure S1A) displays a prominent elution peak at 11.3 ml, corresponding to the channel tetramer (Figure S1B-I). While the R162W mutant of Kir7.1 has more functional disease relevance due to its association with snowflake vitreal degeneration [6], we were unable to extract the R162W mutant protein from the membrane, so instead, focused on a related mutation, R162Q, identified in retinitis pigmentosa patients with less severe disease outcomes [7]. Mutation of Arg162 is implicated in disruption of the PIP_2_ binding site [13, 14]. E276A, another retinitis pigmentosa-associated mutation, is located at the cytosolic G-loop gate of the channel. The E276A migrates to the same position as the wild-type channel. However, the profile of the R162Q mutant is left-shifted (Figure S2A). The addition of the channel blocker ML418 to the R162Q mutant reverted the elution profile of the R162Q to migrate to the same position as the wild-type and E276A mutant (Figure S2B-C). This is a possible indication of the discrete conformational preferences of mutant and blocked channels.

### Overall structure of the human Kir7.1 channel and pore constrictions

We determined structures for apo, PIP_2_-bound and ML418-bound wild-type Kir7.1, and apo mutant counterparts, R162Q and E276A, with varying resolutions (Figure 1; Figures S3-4) These represent the first structures determined of the human Kir7.1 channel and reveal that Kir7.1 partitions into the docked and extended conformers previously described for inwardly rectifying potassium channels [13, 14]. The overall structure of the channel is a tetrameric arrangement, with each monomer conforming to the classical fold of the inwardly rectifying potassium channels. A transmembrane domain consisting of two transmembrane helices, M1 and M2, that are linked from the outer M1 helix through the extracellular turret loop regions before descending to the shorter pore-lining helix that then extends a loop region through the selectivity filter ^119^TIGYGYT^124^ back the extracellular side to connect to the inner M2 helix. This arrangement extends both N and C termini to the cytoplasm, with both termini collectively forming the cytosolic domain. The C-terminal domain is composed of beta-sheets arranged in an Ig-like domain fold. The N-terminal peptide region forms a hook-like structure that folds onto the Ig-like domain and continues to the slide helix at the interface of the transmembrane and cytosolic domains (Figure 1). A single disulfide bond is present at the extracellular turret region from Cys99 near the pore helix to Cys131 at the apex of the M2 helix.

**Figure 1.**
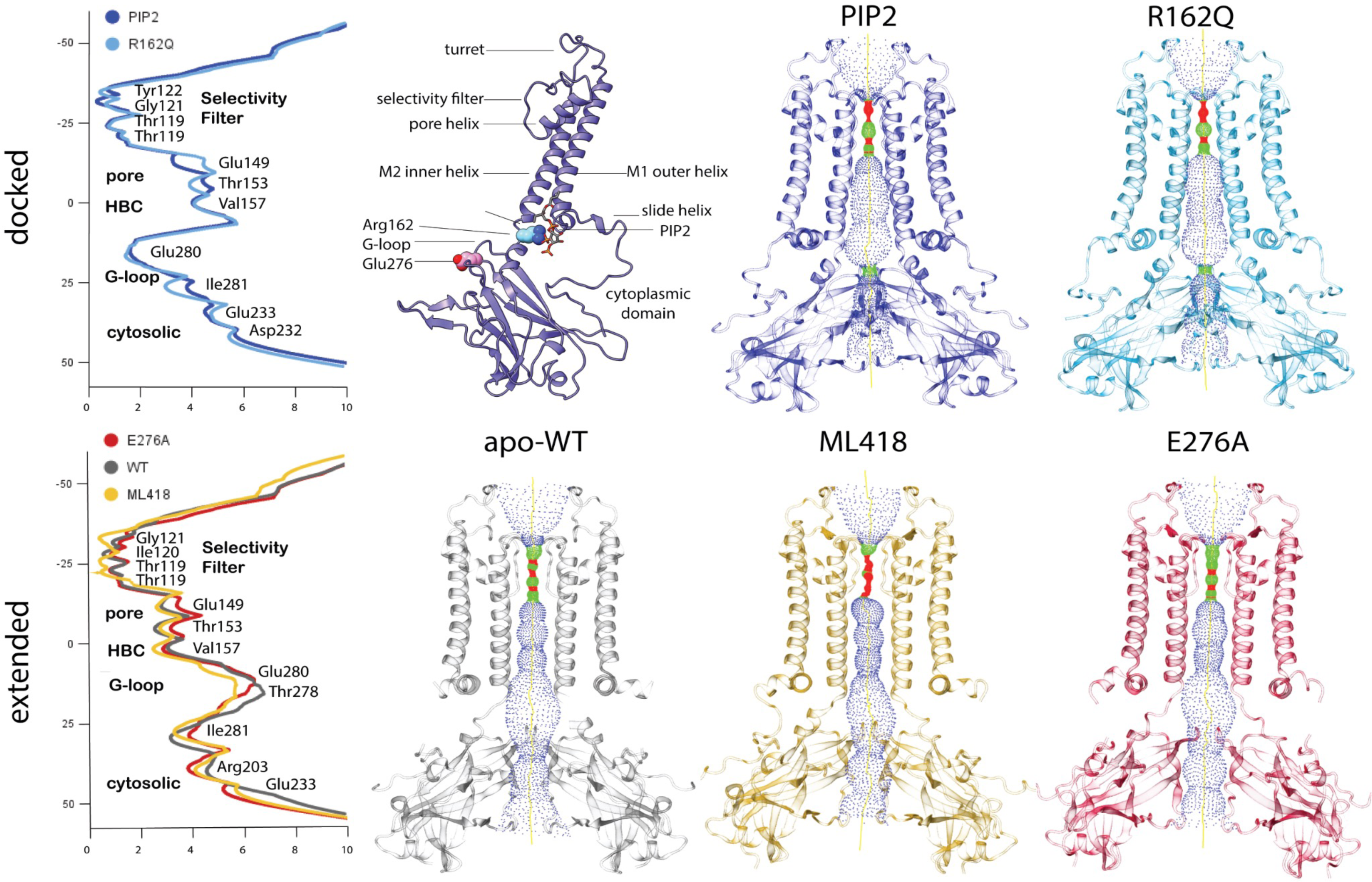
Structure of various forms of the potassium inward rectifier channel Kir7.1. Left. Pore radius calculated between van der Waals surfaces by the program HOLE and plotted as a function of the distance along the pore axis for open conducting docked conformations of PIP_2_ bound Kir7.1 and R162Q mutant (upper left) or for the extended non-conducting conformation of apo WT, ML418 bound and E276A mutant Kir7.1 (lower left). Identities of residues at the narrowest and widest radii along the pore axis are indicated along with associated structural elements. Upper left middle: Ribbon diagram of a single subunit of Kir7.1 tetramer bound to PIP2. Structural elements are labeled, and the location of the PIP2 binding site and retinitis pigmentosa-associated mutations, R162Q and E276A, are indicated. Upper right: Pore volume shown within structures of PIP2 bound WT, R162Q mutant, and for apo WT, ML418 bound and E276A mutant Kir7.1 (lower right). The central pore region is shown as a dotted surface where red is where the pore radius is too narrow for a water molecule, green is where there is room for a single water, and blue is where the radius is double the minimum for a single water molecule.

Using the software HOLE2, we measured the radius between van der Waals surfaces within the ion conduction pathway for each of the structures obtained. Within the compact PIP_2_ bound wild-type and apo-R162Q mutant docked conformation structures, two narrow constriction points are evident and are located at the selectivity filter (Figure 1) and at the G-loop gate, with the narrowest constriction of the G-loop gate occurring at residue Glu280 (Figure 1). In the ML418 bound wild-type, apo-wild-type and apo-E276A mutant structures, constriction is limited to the selectivity filter. While the occupied selectivity filters of the PIP_2_ bound and mutant structures constrict to ∼0.8 Å radius at the narrowest point, the selectivity filter narrows to ∼0.3 Å in the unoccupied ML418 structure (Figure 1). At the G-loop gate, the docked structures are constricted to a 1.0 Å radius in the PIP_2_ bound and 1.2 Å in the R162Q mutant structures. In the extended conformations, the G-loop widens to a radius of 2.6 Å in the E276A mutant and 2.9 Å in the ML418 bound structure (Figure 1). This widening of the G-loop gate is also coincident with a loss of electronegativity and a gain of hydrophobic character at the constriction point. Rather than Glu280 lining the pore as in the docked structures in the extended conformation, a change in the interaction network of residues in the G-loop leads to withdrawal of Glu280 sidechain and protrusion of Ile281 into the pore, thus increasing both the width and hydrophobic character of the constriction at the G-loop gate.

While not as pronounced as the constrictions at the G-loop gate and the selectivity filter, there is a narrowing of the pore at two other locations in the extended conformation relative to the docked conformation. The first occurs at the position equivalent to the helix bundle crossing and occupied by Val157 (Figure 1, 4). Finally, at the exit on the cytosolic side, another constriction appears relative to the docked conformation (Figure 1, 4). In the extended conformer, Arg305 lines the narrowest part of the pore at the cytosolic side, replacing Glu233 seen in the docked conformation. This constriction also increases the electropositive character of the pore at this point and possibly acts in a repulsive way to close the pore.

### Structure of the human Kir7.1 channel in the presence of PIP_2_

Kir7.1 has been shown to be gated by the membrane phospholipid PIP_2_ [15]. To obtain a structure of the open conducting conformation of Kir7.1, the addition of 1 mM C8 PIP_2_ was incubated with preparations of wild-type Kir7.1 prior to deposition on cryo-EM grids. An average resolution of 3.97 Å and local resolution reaching 2.8 Å in the selectivity filter was obtained by cryo-electron microscopy (Figure 1 & Figure S1). As has been observed for Kir2.2 and Kir3.2, the binding of PIP_2_ induces a rotation and anchoring of the cytosolic domain to the transmembrane region, resulting in a compact form of the channel with the dimensions 82 Å by 108 Å. A unique feature of the Kir7.1 channel is the constriction of the G-loop in the open conducting conformation. Rather than a hydrophobic residue, as is present in all other eukaryotic Kir channels, a glutamate residue lines the pore at the G-loop. The distance between glutamate carboxyl groups across the pore is ∼5.4 Å, approximately twice the K-O bond distance in a hydrated potassium ion. This would suggest that at the G-loop in Kir7.1, the passage of potassium proceeds through the replacement of the solvation shell by the carboxyl groups of the glutamate side chain (Figure 1, 3).

### Conservation of the PIP_2_ binding site

The PIP_2_ binding site is highly conserved with minor differences between Kir7.1 and that of those already determined, Kir2.2 and Kir3.2 [13, 14]. The inositol head group of PIP_2_ is coordinated by the residues His26 from the N-terminus, Arg52, Trp53, and Arg54 at the base of the M1 helix adjacent to the slide helix, Met57 from the M1 helix and Lys159, Arg162, and Lys164 from the tether (Figure 2). The P1 phosphate of the inositol headgroup is coordinated by the Arg52 and Arg54 side-chains which are part of the conserved (K/R)WR motif from the outer M1 transmembrane helix. The tryptophan sidechain from this motif is oriented on the opposite side of the helical turn and, along with Met57, co-ordinates the acyl chains from PIP_2_. The P5 phosphate is coordinated by the side chains of Lys159, Arg162, and Lys164 from the tether helix. The basic nature of the side chains in the binding site is preserved between family members with only conservative amino acid substitutions. In Kir3.2, Arg162 is exchanged for a glutamine residue. The lysine at position 159 is invariant between all Kir channels; however, in both Kir2.2 and Kir3.2, the phosphate is additionally coordinated by two vicinal lysines rather than the single lysine, Lys164, in Kir7.1. The P4 phosphate is coordinated by His26 from the N-terminal loop, whereas in Kir3.2, this is replaced by a lysine residue. Determination of the PIP_2_ bound conformation confirmed that the Arg162 residue constitutes part of the PIP_2_ binding site, providing a partial explanation of the occurrence of retinal disease with missense mutations of this residue.

**Figure 2.**
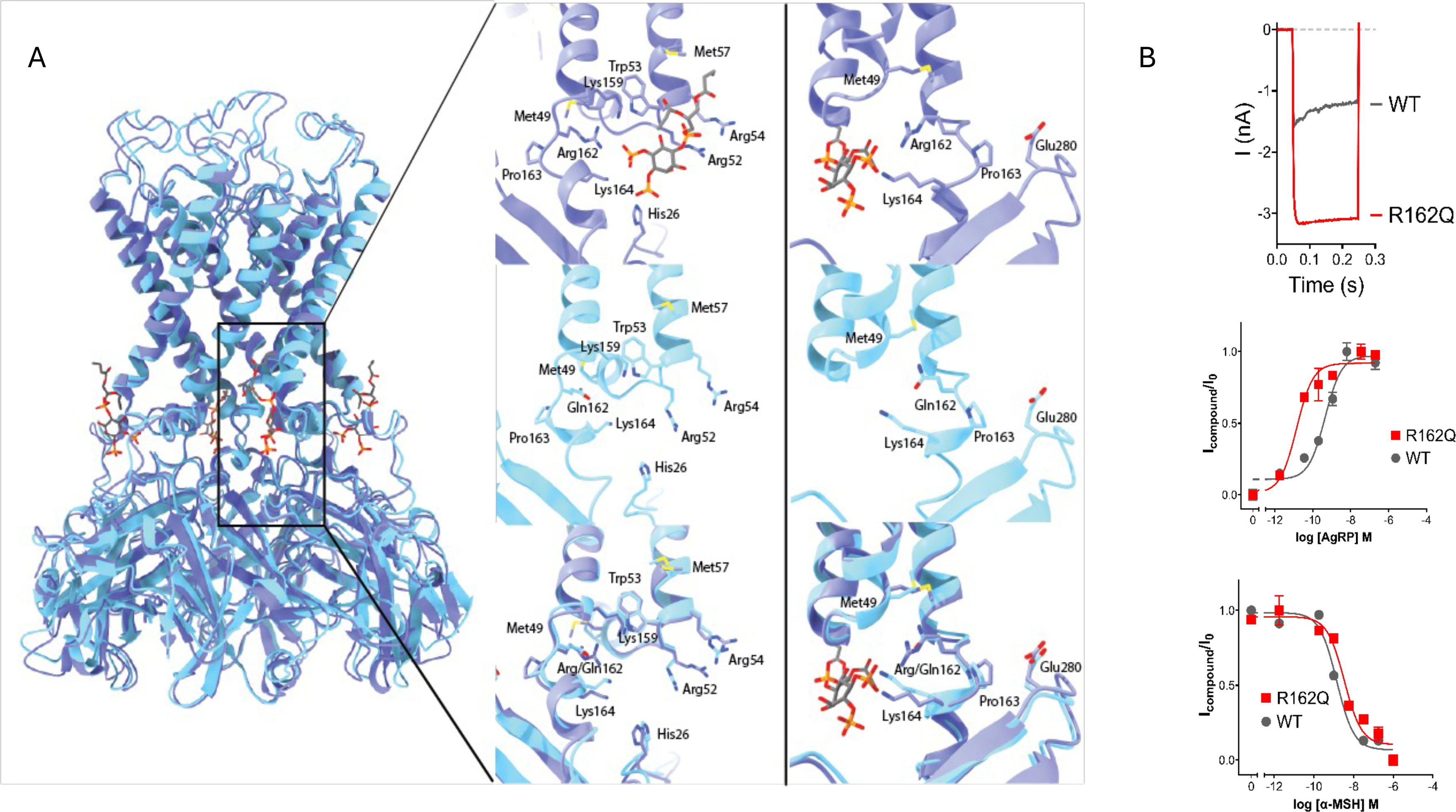
Structure and mechanism of the PIP_2_ bound state and R162Q mutation. (A) Left panel: Alignment of R162Q (blue) mutant atomic model to PIP_2_ bound WT Kir7.1 (purple). All atom RMSD of 1.234 (0.909 among 275 of 306 pairs). Middle panel: PIP_2_ binding site in WT Kir7.1 shown from the front (upper middle), view of transmembrane regions showing arrangement of PIP_2_ binding residues in R162Q mutant structure (middle) and overlay of PIP_2_ bound WT and R162Q (lower middle). Right panel: Configuration of C-linker in PIP_2_ bound WT (upper right) and in R162Q mutant (middle right) displaying residues important for maintaining helicity of C-linker and overlay of PIP_2_ bound WT Kir7.1 with R162Q mutant (lower right). The presence of glutamine over arginine allows the channel to adopt the open conformation in the absence of PIP_2_ by substituting for the P5 phosphate, allowing for a bridging interaction between Lys164and Gln162 and Gln162 to Met49 to dock C-linker to TM1. The G-loop interaction of Pro163 from the linker to Glu280 from the G-loop is maintained. (B) Top: representative current traces for HEK293 cells expressing either WT or R162Q Kir7.1, from a holding potential of −60 mV to a step pulse of −140 applied for 200 ms. Bottom: Concentration response curves in cells with co-expression of the MC4R and either WT or the R162Q mutant Kir7.1, showing the effects of AgRP or α-MSH on the functional coupling of MC4R to the Kir7.1 channel.

### R162Q mutant adopts near identical conformation as PIP_2_ bound wild-type channel

We found that the introduction of the retinitis pigmentosa-associated R162Q mutation into Kir7.1 induced a conformational stabilization (Figure 2A), enabling us to obtain reconstructions of 3.5 Å resolution with local resolution approaching 2.8 Å. Alignment of Cα atoms of the mutant protein to the PIP_2_ bound wild-type channel resulted in a global all-atom RMSD of 1.234 and an RMSD of 0.909 among 275 best matching pairs (Figure 2A). Overall, the configuration of residues in the PIP_2_ binding site is conserved and although very nearly identical, there are some notable differences between the two structures. When mutated to glutamine, residue 162 coordinates with the Met49 in the slide helix near the base of the outer M1 transmembrane helix within the same subunit. This glutamine residue in the R162Q mutant effectively substitutes for the P5 phosphate and to maintain the helicity of the interdomain tether, thus docking the cytosolic domain to the transmembrane region (Figure 2). In the PIP_2_ docked structure, Met49 forms a hydrophobic interaction with Thr278 from the G-loop in an adjacent subunit. Docking of the cytosolic domain to the transmembrane region in the PIP_2_ bound structure is achieved predominantly through interactions of the slide helix at the transmembrane interface with the G-loop and CD-loop on the cytosolic domain. In the R162Q mutant structure these docking interactions are mostly through the slide helix interaction with the tether helix. Between the two structures, the interaction of Asp43 from the slide helix with Asn165 from the tether is preserved. In addition, interactions between M1 and M2 transmembrane helices are altered in the R162Q mutant, with Met57 free to interact with Met147 across the helices. In the PIP_2_ bound structure, Met57 is engaged with PIP2, leaving Met147 to stack between Phe156 in the M2 helix and Phe60 in the M1 helix in an adjacent subunit. These differences may account for subtle differences in the pore dimensions (Figure 1).

We also characterized the functional properties of the R162Q mutant by transfecting the mutant Kir7.1 channel alone or the channel plus the WT MC4R into HEK293 cells, followed by whole cell patch clamp recordings. Recordings show that Kir7.1 R162Q affected ion channel permeation. Cells expressing the Kir7.1 R162Q mutation exhibit a gain-of-function phenotype. In whole-cell patch clamp recordings, the R162Q mutant channel shows increased potassium currents compared to the wild-type channel (Figures 2B and S7). Further, we determined that the Kir7.1 R162Q mutation affected the functional coupling between MC4R and Kir7.1. While the mutation did not impact the EC_50_ of channel closure by α-MSH, it did significantly increase the EC_50_ of channel opening by AgRP (Figures 2B and S7).

### E276A mutant adopts an extended conformation

In apo wild-type and E276A structures, Kir7.1 is observed in the extended conformation in a manner equivalent to that seen in the unliganded cryo-EM structures of GIRK2 (Figure 3). Cytosolic and transmembrane domains are disengaged, and contacts between the transmembrane region and the G-loop and tether helix are lost. This extension is achieved by an unraveling of the tether helix between the two domains, with the distance between the domains increasing by an almost equivalent rise of a single helical turn from 7.2 Å to 11.6 Å and through rotation of ∼24° by the cytosolic domain relative to the PIP2 bound structure (Figure 3) when aligned through the transmembrane regions.

**Figure 3.**
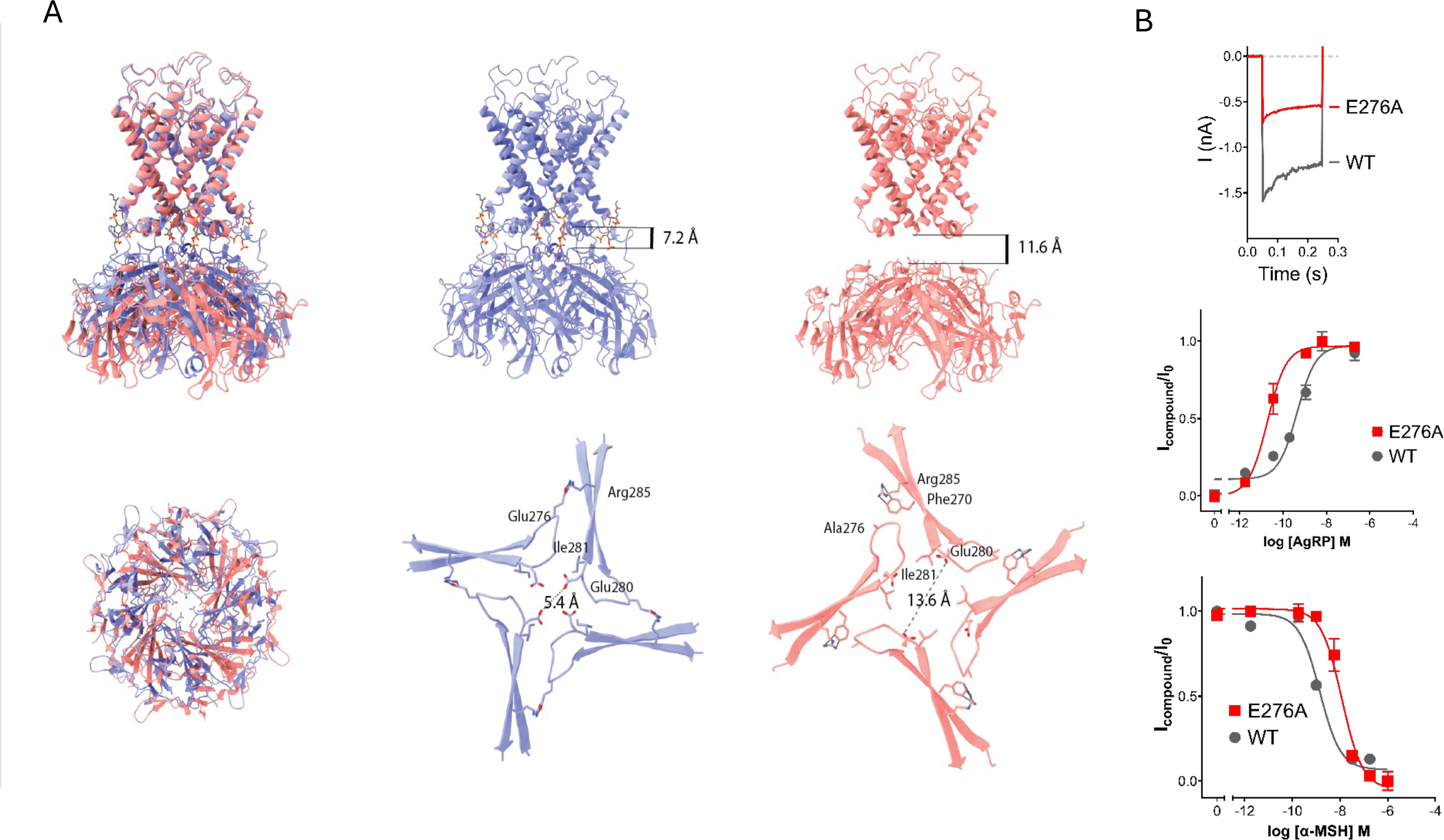
Structure and mechanism of the E276A mutation. (A) PIP_2_ bound wt Kir7.1 (purple) aligned by transmembrane domains to the E276A mutant (salmon). Upper left: side view of the entire channel. Lower left: cytosolic domain as viewed from the transmembrane side above the cytosolic G-loop. Upper middle and right. Ribbon diagram of PIP_2_ bound WT and E276A mutant with distances between transmembrane and cytosolic domains indicated. Increased distance between the transmembrane and cytosolic domain in the extended conformation due to linker helix unraveling and disengagement of cytosolic domain with slide helix. Lower middle and right: View of G-loop displaying placement of residues Glu280 and Ile281 as viewed down the central pore. Distance from diametrically opposed Glu280 carbonyls displayed. (B) Top: representative current traces for HEK293 cells expressing either WT or E276A Kir7.1, from a holding potential of −60 mV to a step pulse of −140 applied for 200 ms. Bottom: Concentration response curves in cells with co-expression of the MC4R and either WT or the E276A mutant Kir7.1, showing the effects of AgRP or α-MSH on the functional coupling of MC4R to the Kir7.1 channel.

In wild-type Kir7.1, Glu276 is located at the interface between subunits within the channel tetramer, with Glu276 from one chain coordinating Arg285 from the adjacent chain in the wild-type structure. We found that the E276A mutant was preferentially stabilized in the extended conformation (Figures 1, 3) and were able to obtain <5 Å reconstructions (Figure S3). When Glu276 is mutated to alanine, the contact with Arg285 from an adjacent subunit can no longer be formed, and Arg285 instead folds back to engage in pi-cation interactions with Phe236 in an adjacent subunit (Figure 3). The result is a loss of the constriction point in the channel at the G-loop gate (Figure 3), with the G-loop widening and becoming more hydrophobic in character lined by Ile281 rather than Glu280. Interestingly, Phe236 occupies an equivalent position as one of the sodium binding residues in the GIRK structures.

We also characterized the functional properties of the E276A mutant, as described above. Recordings show that Kir7.1 E276A also affected ion channel permeation. However, in contrast to the R162Q mutant, the E276A mutant channel shows reduced potassium currents compared to the wild-type channel (Figures 3B and S7). Further, when co-expressed with the MC4R, this mutation impacts both α-MSH and AgRP-mediated channel function, increasing sensitivity to AgRP-mediated channel opening and decreasing sensitivity to α-MSH-induced channel closure.

### Channel blocker ML418 resides in the inner vestibule beneath the selectivity filter

Pharmacological studies have supported the hypothesis that the compound ML418 is a blocker of the Kir7.1 channel [12]. In the presence of the channel blocker ML418, Kir7.1 was observed to assume an extended conformation (Figure 4A). Although lower resolution reconstructions were obtained ∼4.95 Å (Figure S4), we were able to model the protein into the density and make predictions about interactions with the channel blocker and the residues involved in conformational changes. ML418 enters from the intracellular side of the pore and is located at the upper limits of the pore cavity extending towards the S4 site of the selectivity filter (Figure 4A). Sufficient density was present to place piperidine and quinoline rings within the channel vestibule. The isopropyl carbamate group of ML418 was refined to be present beneath the selectivity filter with the ester oxygens coordinated by Thr119. Although the S4 site is typically occupied by potassium in two separate structures of KirBac [16, 17], a magnesium or calcium ion is observed at this position, while the modeled piperidine ring occupies the equivalent position to strontium in the human Kir2.1 structure [18]. Strong residual density from one Glu149 in one subunit of Kir7.1 from the M2 helix is observed potentially due to the coordination of the piperidine nitrogen of ML418. Additional coordination may be provided by Thr153 in the pore cavity (Figure 4). One of the greatest changes is seen at the selectivity filter, which appears to be unoccupied and in a collapsed state in the ML418 bound structure. There are also differences in the interaction network between the transmembrane helices that may allow for the widening of the pore at the cytosolic gate. In the PIP_2_ bound structure, Val157 is drawn away from the pore cavity in the PIP_2,_ while in the ML418 bound structure, it occupies the pore (Figure 4). Due to the difficulty of accurately placing an asymmetric ligand within a symmetric assembly, we attempted refinement without applying symmetry. While this resulted in a loss of resolution, some interesting features emerged with altered inter-subunit contacts evident within the channel. Differences were observed within the turret region, TM1 helix, and at the slide helix. Particularly notable is the continuous density between intersubunit TM1 and the slide helix, suggesting a contact between Trp45 in the slide helix and Trp53 at the base of TM1. This contact would occlude the PIP_2_ binding site and suggest a potential desensitized state of the channel. The alteration of contacts with turret, pore helix, and transmembrane regions suggests that ML418 occupancy of the selectivity filter and central pore region propagates changes in TM1 that allow contact between neighboring subunits (Figure 4S). As reported previously [12], ML418 potently inhibits conductance through the Kir7.1 channel (Figures 4B and S8), and ML418 sensitivity is not affected by co-expression of MC4R with Kir7.1 8 (Figures 4B and S8).

**Figure 4.**
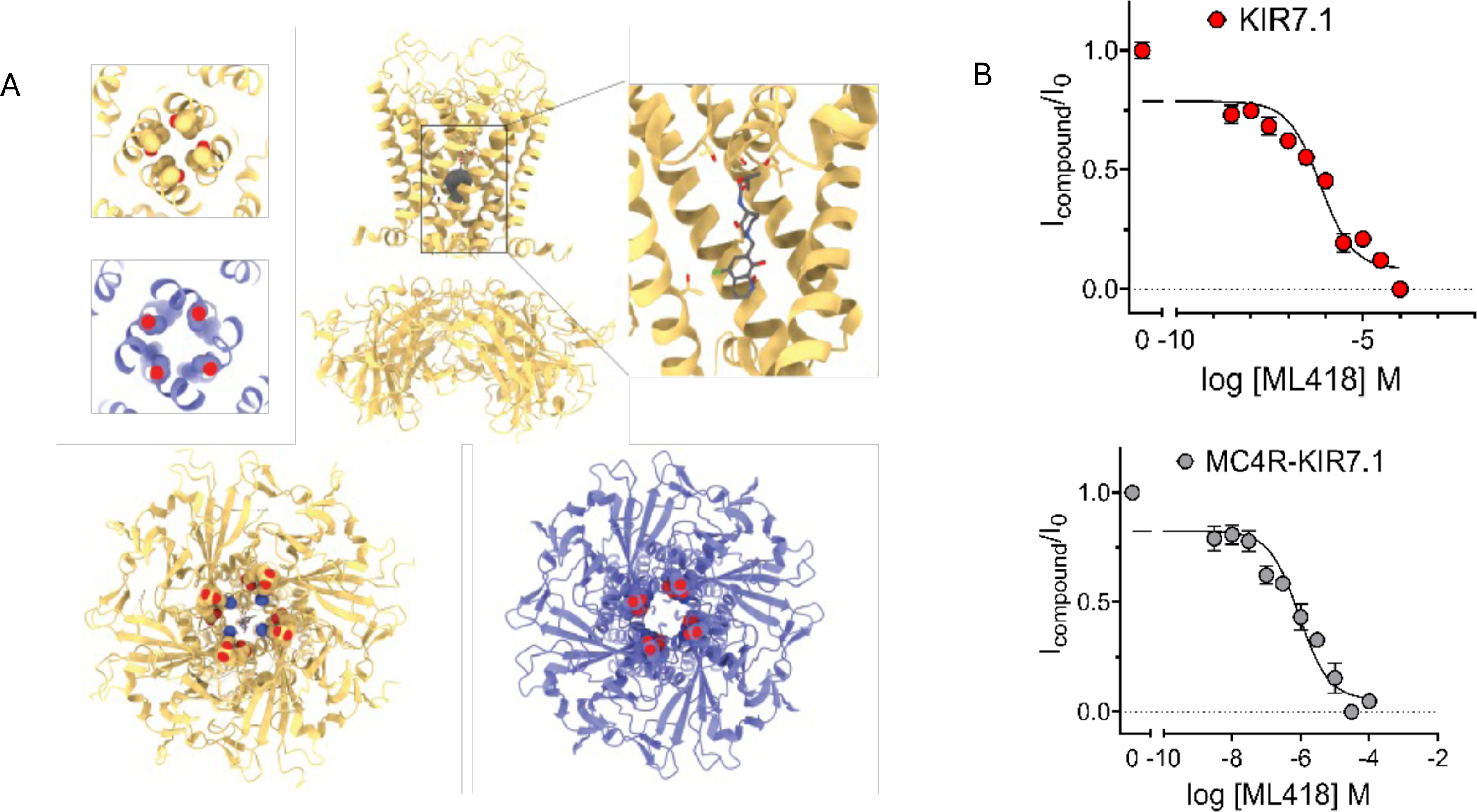
Structure and mechanism of the channel blocker ML418. (A) Upper left: View of helix bundle crossing in ML418 bound (yellow) and PIP_2_ bound WT Kir7.1 (purple) showing restriction in non-conducting conformation of ML418 bound structure. Upper middle: Ribbon diagram of ML418 bound Kir7.1 refined in C4 symmetry with density due to ML418 displayed in the central pore. Upper right: refined position of ML418 in Kir7.1 central cavity. Lower panel: view of the cytosolic exit side of ML418 bound Kir7.1 (left, yellow) and PIP_2_ bound Kir7.1 (right, purple) displaying altered charge profile of pore-lining residues. (B) Concentration response curves for ML418 in cells with expression of Kir7.1 alone or Kir7.1 plus MC4R.

### Inhibition of Kir7.1 decreases food intake and promotes weight loss

With structural and pharmacological data supporting ML418 as a blocker of the Kir7.1 channel and data suggesting that α-MSH depolarizes and activates MC4R neurons by blocking Kir7.1 [9], we next used this compound to test the *in vivo* activities of Kir7.1 in the regulation of feeding behavior and body weight. We administered ML418 (from 5 to 7.5 mg/kg) or vehicle peripherally to wild-type mice (Figure 5A). ML418 decreased 24-hour food intake and induced 24-hour weight loss in a dose-dependent manner (Figure 5B-C), without impacting water intake (Figure 5D). ML418 also has a high affinity to SUR1/Kir6.2 channels (K_ATP_) [12]. To test if the anorectic activity of ML418 was due to K_ATP_ channel inhibition, we administered A-4166 (nateglinide) (5 mg/kg), a potent Kir6.2 inhibitor [19], to wild-type mice. A-4166 did not affect acute feeding in mice (Figure 5E). Kir7.1 is widely expressed, and global deletion of Kir7.1 leads to postnatal death due to improper development of the lungs and palate [20]. Thus, we also tested whether central administration of ML418 affected feeding and body weight (Figure 5F). Lean cannulated mice received ML418 or vehicle. We found that ML418 delivered centrally significantly reduced food intake (Figure 5G), whereas a Kir4.1 inhibitor, VU0134992 [21], did not affect food intake (Figure 5H). Next, we tested the potential of ML418 to decrease food intake in DIO mice with increased adiposity. Remarkably, a single central administration of ML418 induced a sustained reduction in both food intake and weight over 5 days (Figure 5J-K).

**Figure 5.**
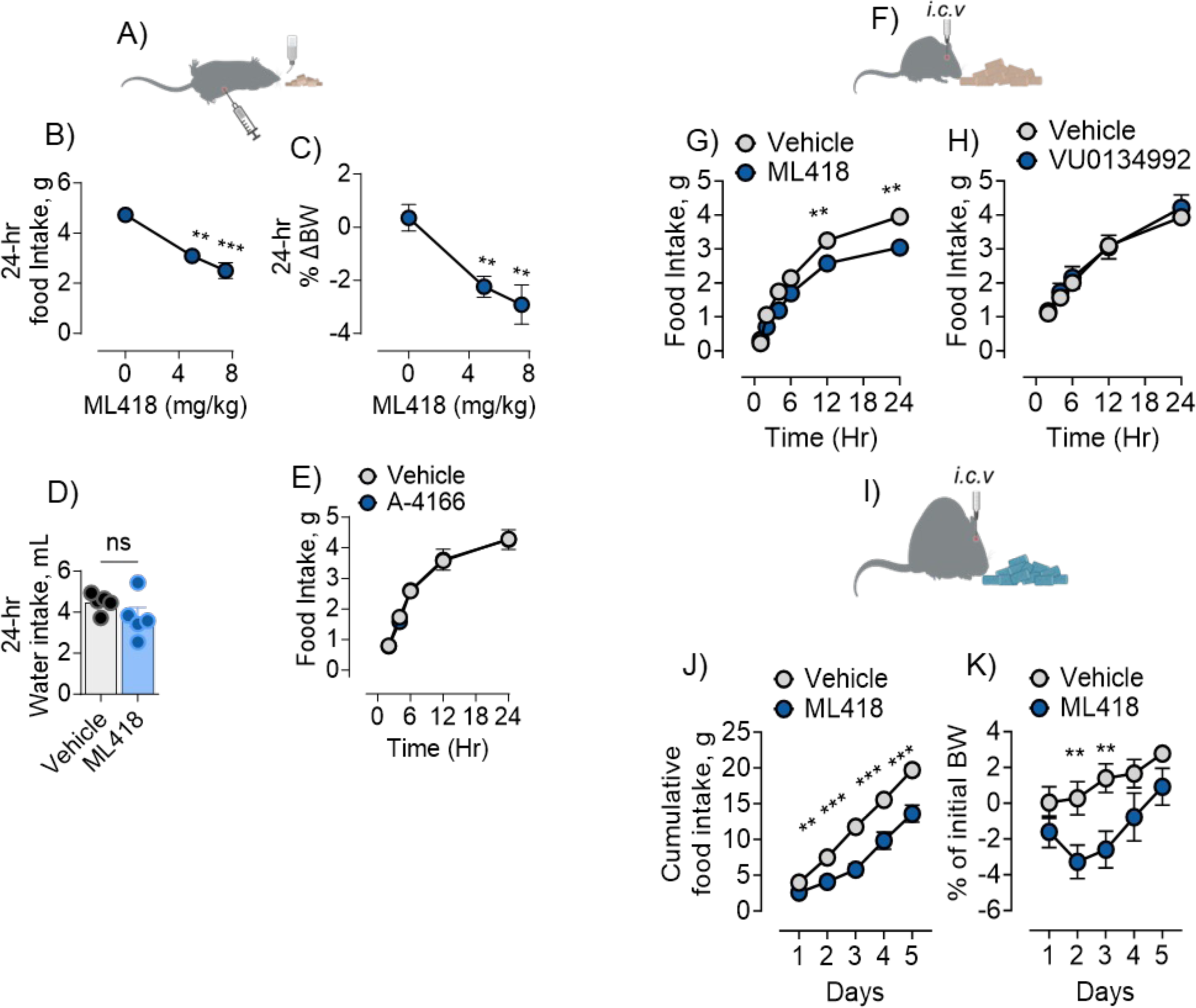
Inhibition of Kir7.1 decreases food intake and promotes weight loss independent of adiposity. (A) Depiction of experimental mice fed standard chow receiving intraperitoneal injection (ip). (B) Food intake of wild-type C57BL/6J mice over 24-hour period treated with 13% DMSO in saline (vehicle) or ML418 (5 mg/kg & 7.5 mg/kg); n= 7-13, one-way ANOVA, F (2, 23) = 21.87, p<0.0001. (C) 24-hour change in body weight in wild-type mice treated with 13% DMSO in saline (vehicle) or ML418 (5 mg/kg & 7.5 mg/kg); n= 7-13, one-way ANOVA, F (2, 22) = 9.988, p<0.001. (D) Water intake of wild-type mice over a 24-hour period treated with vehicle (13% DMSO in saline) or ML418 (7.5 mg/kg), n =5, Student’s t-test (2-tailed), t(8) = 1.361, p>0.05 (ns). (E). Food intake in wild-type mice treated with A-K4166 (5 mg/kg) or vehicle (13% DMSO in saline), n =5, two-way ANOVA, F (4, 40) = 0.06315, p>0.05. (F) Depiction of mice receiving intracerebroventricular injection (icv) with free access to standard chow diet. (G) Acute food intake over 24-hours in wild-type mice centrally treated with vehicle (1uL DMSO) or ML418 (500picomoles, 1uL), n=7-9, two-way ANOVA, F (6, 98) = 2.841, p<0.01. (H) Acute food intake in wild-type mice centrally treated with vehicle (1uL of DMSO) and VU0134992 (500pmoles/1uL), n =5, F (5, 38) = 0.07213, p>0.05. (I) Depiction of experimental diet-induced obese (DIO) wild-type mice with free access to 60% high-fat diet. (J) Food intake of DIO wildtype mice receiving a single icv injection of vehicle (1uL of DMSO) or ML418 (500pmoles, 1uL), n =9, two-way ANOVA, F (4, 80) = 3.933, p<0.01. (K) Change in body weight of DIO mice receiving single icv injection of vehicle or ML418, n = 9, two-way ANOVA, F (4, 80) = 0.6236, p>0.05. ANOVA tests were corrected for multiple comparisons using the Tukey–Kramer method. Mean ± SEM is shown.

### ML418 activates PVH MC4R neurons, and Kir7.1 in MC4R neurons mediates effects of ML418 on food intake

Building upon our observation that inhibition of Kir7.1 by channel blocker ML418 reduces food intake and promotes weight loss, we next wished to determine if Kir7.1 in MC4R neurons might be mediating the effects of ML418. Based on our earlier observations that α-melanocyte-stimulating hormone (α-MSH) depolarizes PVH MC4R neurons [9], we investigated the impact of ML418 on the depolarization of MC4R neurons by α-MSH. Using whole-cell patch recordings of PVH MC4R neurons from hypothalamic slices derived from MC4R-GFP mice, we reproduced the finding that α-MSH reliably depolarized PVH MC4R neurons (Figure 6A). Treatment with ML418 alone did not alter the baseline membrane potential of the MC4R neurons (Figure 6B). However, ML418 markedly blocked the α-MSH-induced depolarization of PVH MC4R neurons (Figure 6B). Given that administration of ML418 reduces feeding in the mice, we tested whether ML418 activated PVH neurons. Using c-Fos as a marker of neuronal activation, we found a significant increase in c-Fos expression in the PVH upon ML418 treatment (Figure 6C-E). Approximately 70 % of MC4R neurons were c-Fos positive after ML418 treatment, compared to fewer than 20 % following vehicle treatment, indicating that inhibiting Kir7.1 activates the majority of MC4R neurons, along with a significant number of other unidentified PVH neurons.

**Figure 6.**
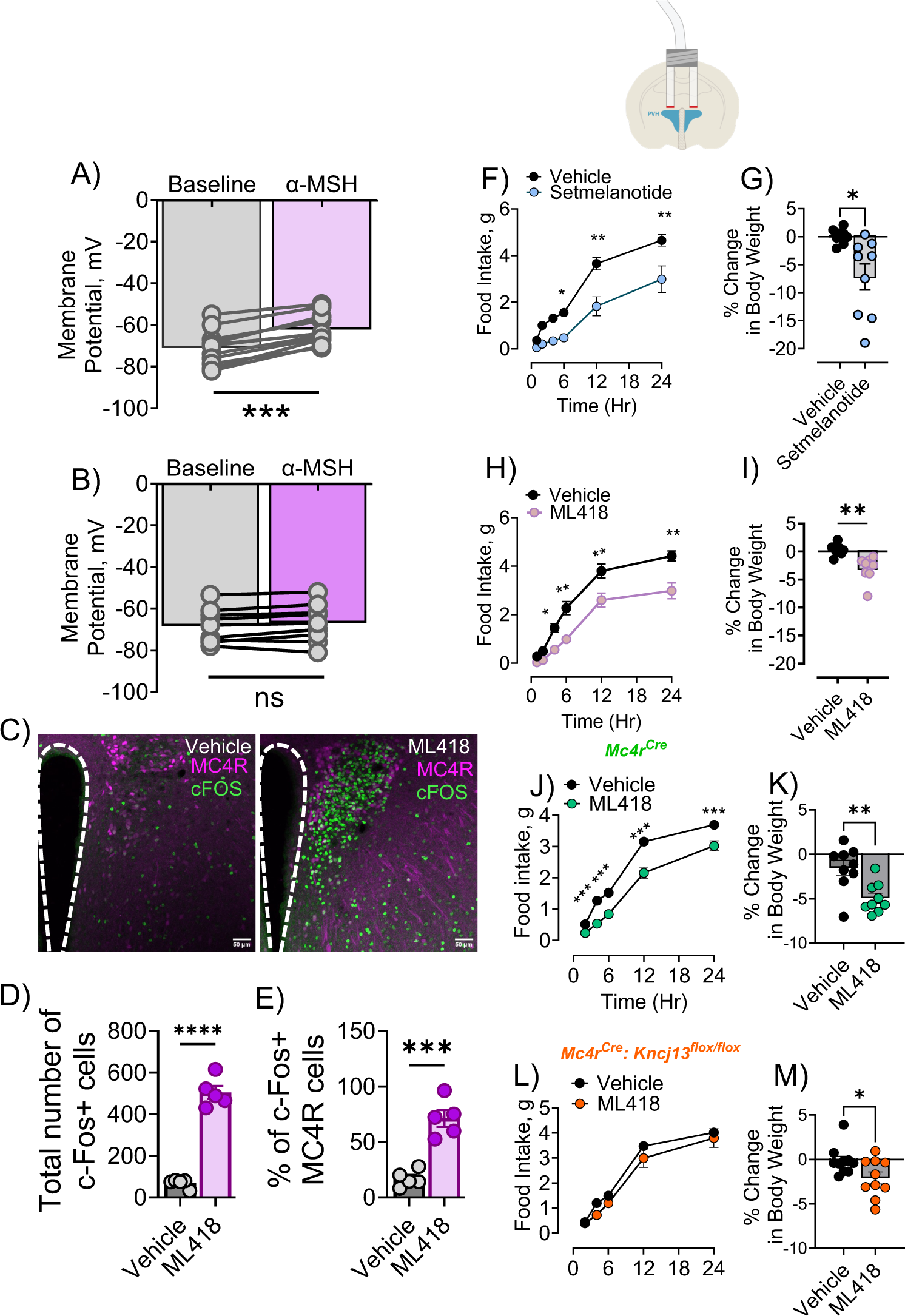
ML418 activates PVH MC4R neurons, and deletion of *Kcnj13* from *Mc4r*-expressing cells blunts feeding behavior. (A) Effect of α-MSH (250 nM) on the membrane potential of PVN MC4R neurons, n= 12 cells. (B) Effect of α-MSH on the membrane potential of PVH MC4R neurons after incubating slices with ML418 (5µm), n= 9 cells. (C) Representative images of coronal sections of the paraventricular hypothalamus (PVH) from MC4R-GFP mice systemically stimulated with vehicle (DMSO) or ML418. (D) Quantitative analysis of cells expressing cFos in the PVH after animals were treated with vehicle or ML418, n= 5, Student’s t-test (2-tailed), t(8) = 13.26, p<0.0001. (E) Percent of MC4R cells that are c-Fos-positive, n =5, t(8) = 6.6109, p<0.001. (F) Food intake in intra-PVH cannulated wild-type mice receiving central injection of vehicle (0.5uL/side, Ringer’s solution) or setmelanotide (250pmoles/0.5uL/side), n=7-9 (5, 84) = 2.258, p=0.0559. (G) 24-hour body weight change of the same mice, Student’s t-test (2-tailed), t(15)=2.908, p<0.05. (H) Intra-PVH cannulated wild-type mice receiving central injection of vehicle (0.5uL/side, Ringer’s solution) or ML418 (250pmoles/0.5uL/side), n=8 F (5, 84) = 3.09,p<0.05. (I) 24-hour body weight change of the same mice, Student’s t-test (2-tailed), t(15)=3.807, p<0.01. (J) Food intake over a 24-hour period of control *Mc4r-Cre* mice treated with vehicle (13% DMSO in saline, ip) or ML418 (5mg/kg, ip), n =9, F (4, 80) = 2.212, p<0.05 and (K) change in 24-hour body weight in the same mice, Student’s t-test (2-tailed), t(16)=3.402, p<0.05. (L) Food intake over 24-hour period of *Mc4r-Cre::Kcnj13^flox/flox^* mice treated with vehicle (13% DMSO in saline, ip) or ML418 (5mg/kg, ip), n =9, F (4, 90) = 0.4581, p>0.05 and (M) change in 24-hour body weight in the same mice, Student’s t-test (2-tailed), t(18)=2.286, p<0.05. ANOVA was corrected for multiple comparisons using the Tukey–Kramer method. Mean ± SEM is shown.

As expected, the injection of setmelanotide (Imcivree) [22], an MC4R agonist, into the PVH led to decreased food intake and weight loss (Figure 6F-G). Direct administration of ML418 into the PVH of wild-type mice (Figure 6D) similarly resulted in reduced food intake and weight loss within 24 hours (Figure 6H-I). To determine if Kir7.1 in MC4R neurons plays a role in inhibition of feeding and weight loss following ip administration of ML418, we used the Cre-LoxP system to delete *Kcnj13*, the gene encoding Kir7.1, from MC4R neurons. As expected, *Mc4r-Cre* mice exhibited a significant reduction in food intake and body weight following ip ML418 injection (Figure 6J-K). Conversely, mice with selective deletion of *Kcnj13* from *Mc4r*-expressing cells had no changes in food intake following ip ML418 injection and exhibited less weight loss relative to ML418 treated control *Mc4r-Cre* animals (Figure 6L-M). Interestingly, no discernible effect on acute food intake or body weight was observed upon conditional deletion of *Kcnj13* from *Mc4r*-expressing cells, suggesting potential compensatory mechanisms during development.

### Testing the structure and function of an MC4R:Kir7.1 complex

The data described above provides strong support for the role of Kir7.1 in α-MSH- and AgRP-mediated regulation of feeding behavior and body weight via the MC4R. Further, pharmacological data shows that G protein signaling is not required for MC4R-mediated regulation of Kir7.1 [9]. To explore the possibility of complex formation between MC4R and Kir7.1 as requisite for the direct gating of the channel by melanocortin ligands, we pursued purification of the receptor and channel as a single polypeptide chain. Towards this end, the MC4R coding sequence was fused at the carboxy-terminus to the amino-terminus of the Kir7.1 coding sequence with no linker or with linkers containing 20, 30, and 40 amino acids (Figure 7). Except for the construct containing no linker, the fusion of Kir7.1 to MC4R with linkers L1-L3 had no impact on the EC_50_ of α-MSH-mediated MC4R coupling to Gs, as monitored using an assay for intracellular cAMP (Figure 7A-C). While constructs L_0_, L1, and L2 impaired the maximal activity of the MC4R, construct L3 exhibited a normal EC_50_ and 100% Emax relative to the WT receptor (Figure 7). While the tethered constructs exhibited reduced basal conductance, all 4 constructs exhibited a concentration-responsive increase in conductance upon treatment with AgRP, with nanomolar EC_50_ values (Figures 7 and S9).

**Figure 7.**
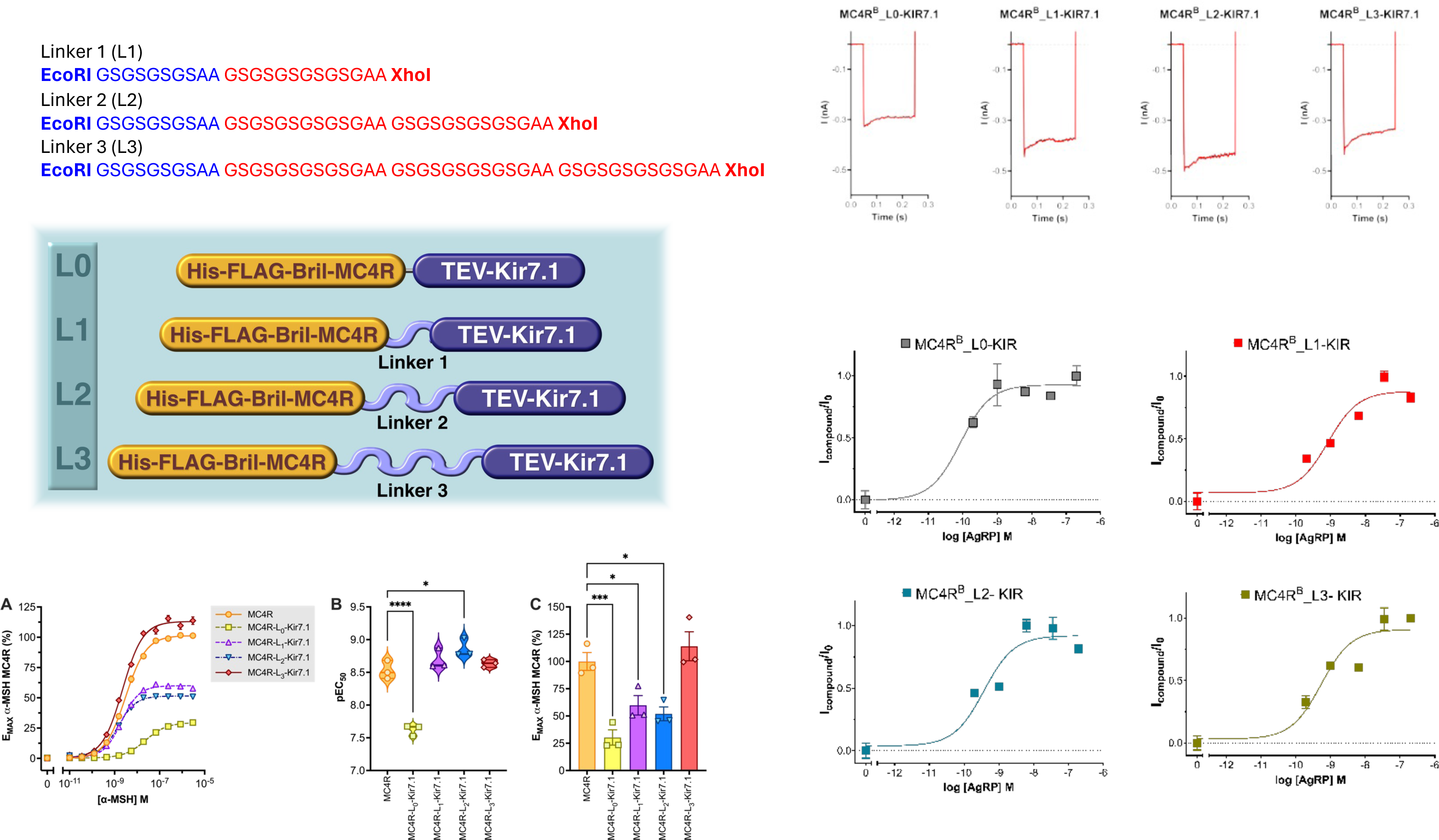
Creation of a linkered MC4r-Kir7.1 protein that retains receptor-mediated channel regulation. Top Left: Structure of linkered MC4R-Kir7.1 constructs with no linker or linkers of indicated sequence (L1 through L3). The bottom left shows (**A**) Intracellular cAMP production concentration-response curves in cells transfected with the GScAMP22f luciferase cAMP genetic sensor and the indicated MC4R and Kir7.1 tethered constructs. The data represent the mean ± S.E.M. from twelve to fifteen replicates distributed in three assay plates. The data were fitted to a four-parameter sigmoid model. (**B**) pEC_50_ values and (**C**) maximum effect (E_MAX_) relative to the control MC4R construct transfection obtained from the concentration-response curves shown in A. The data points represent the pEC_50_ or E_MAX_ values determined from each of the three assay plates. Significance was determined by one-way ANOVA and the Dunnett post-test.* = p<0.05, ** = p<0.01, *** = p<0.001, **** = p<0.0001. Top Right: The representative current traces of the MC4R-Kir7.1 tethered constructs are shown, evoked from a holding potential of −60 mV to a step pulse of −140 applied for 200 ms. Bottom Right: Concentration response curves of AgRP measured from cells expressing the MC4R-Kir7.1 tethered constructs.

In the presence of ML418, we were able to obtain preparations of the MC4R-L2-Kir7.1 complex where a well-resolved peak was evident from the size exclusion profile over a Superose 6 column eluting at ∼15.5 mL (Figure 8A-E). A Western blot for FLAG peptide and Kir7.1 confirmed the presence of both MC4R and Kir7.1 as a single chain with higher bands corresponding to dimer and tetramer also evident. Some degradation bands were also present. We were able to obtain cryo-EM reconstructions using applied C4 symmetry from a very small number of particles. The presence of two conformations appeared, one more compact than the other and likely corresponding to the docked and extended conformations already observed for the isolated channel. Also apparent was an alternate orientation of the external density surrounding the central pore in the reconstructions relative to the axis along the membrane. Using alphafold to predict the complex resulted in a prediction of interactions between the receptor, the transmembrane and the cytosolic domains of the channel. Deeming the cytosolic interactions as unlikely to be accurate, we proceeded with the prediction of the transmembrane interaction only. This provided a reasonable fit to the compact reconstruction likely representing the open conformer of the channel with an “upright” orientation of the receptor relative to the membrane, and by aligning the transmembrane component of the prediction to the open conformation of the channel we generated a model of AgRP bound MC4R complexed with Kir7.1. This suggests interactions principally through MC4R TM5 with the M1 helix of Kir7.1 at the extracellular interface, MC4R ICL2 with the Kir7.1 slide helix and MC4R ICL3 with the cytosolic ß-hairpin and N-terminal peptide are responsible for maintaining the channel open conformation. Starting from the alphafold prediction we then aligned the inactive SHU9119-bound MC4R crystal structure to the alphafold prediction and then active G-protein bound cryo-EM structure of setmelanotide: MC4R was aligned through the peptide binding site, and the docked open conformer was replaced with the extended nonconducting conformer of the Kir7.1 channel aligned through the transmembrane regions. This generated a receptor with a tilt relative to the membrane consistent with that observed for the cryo-EM reconstruction of the more extended conformation. This tilted conformation leads to disengagement of MC4R with the cytosolic domain of Kir7.1 and in an orientation of the receptor relative to the channel compatible with G-protein binding.

**Figure 8.**
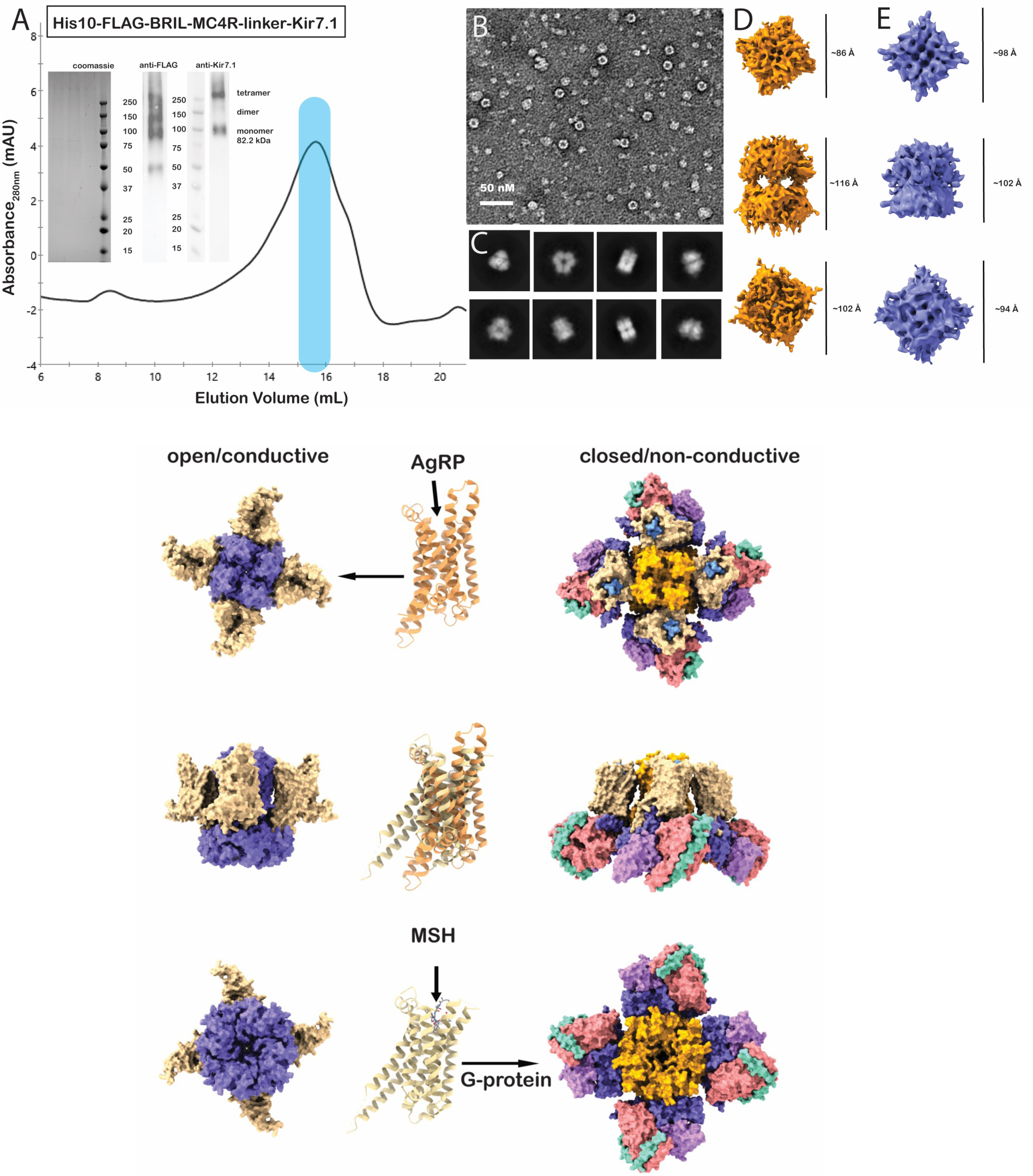
Purification and modeling of a linkered Kir7.1/MC4R structure. Upper left: Size exclusion profile of L2-linked MC4R:Kir7.1 construct. Coomassie-stained and anti-FLAG and anti-Kir7.1 western blot. Middle: negative electron micrograph of peak fraction from size exclusion profile and 2D class averages of cryo electron micrographs. Right: 3D cryo - EM volumes obtained from two major classes with dimensions indicated. Lower: Models of MC4R bound to Kir7.1 obtained from alphafold (left) in the open conducting conformation and of active MC4R bound to G-protein in complex with the ML418 bound conformation of Kir7.1 obtained by fitting SH911 bound structure to alphafold prediction then aligning with active MC4R (7PIU) peptide binding site. Tilted conformation corresponds with outward movement of TM5 and TM6 and disengagement of Kir7.1 cytosolic domains by MC4R to allow for G-protein binding.

## DISCUSSION

To control ion permeation, K^+^ channels must switch between activated, resting, and inactivated states. Ions cross through the membrane at the narrowest part of the channel, the selectivity filter, a process that involves transient loss of water by exchange of the solvation shell for carbonyl groups within the filter. Although some potassium channels conduct through very wide pores, it has been suggested that for the Kir channels, the limiting width for permeation is the ionic diameter of potassium (<3 Å) and that transient loss of water from the hydration shell can accommodate permeation through narrow pore widths. In potassium channels, the allosteric coupling between the activation gate and the selectivity filter has long been studied predominantly in the context of c-type inactivation. Canonically, potassium channel conductance through the selectivity filter is gated by coupling, through collective motion of the channel helices, to a conformational change that alternately dilates and constricts a collar-like intracellular entrance to the pore, the activation gate [10, 11]. In Kir channels, the activation gate is the helix bundle crossing or hydrophobic collar, which opens and closes in response to a variety of stimuli [16, 17]. In the voltage-gated potassium channels, after prolonged opening of the activation gate, the selectivity filter switches from the conductive to a restricted conformation, a process known as c-type inactivation. There is accumulating evidence that the conformational changes governing the gating of K^+^ channel pores are not universal.

Using cryo-EM, we have determined structures of Kir7.1 in conducting and non-conducting states by obtaining data from samples in the presence of the Kir channel opener PIP_2_ [13, 14], and with the small molecule channel blocker ML418 [12], which prevents ion conduction through the pore. We have also determined structures of mutated forms of Kir7.1 that have been associated with retinitis pigmentosa and have altered conductance properties. The mutations occur at two key sites that regulate channel opening: Arg162 within the PIP_2_ binding site and Glu276 at the cytosolic G-loop gate. The mutant proteins were observed to be trapped in different conformational states, R162Q adopting a docked conformation and E276A adopting an extended conformation. Collectively, structures of PIP_2_-, ML418-bound, and mutant structures reveal key transitions along the gating pathway with the R162Q in a docked but non-conductive state and E276A and ML418 structures in a non-conductive extended conformation. In addition, when bound to ML418, the selectivity filter is observed to be unoccupied and in a constricted state. We observed a high degree of conformational heterogeneity in our initial preparations of the wild-type unliganded channel, suggesting that the Kir7.1 channel is intrinsically dynamic and oscillates between intermediate conducting and non-conducting states.

The structures of both the mutant E276A and that of Kir7.1 in the presence of the channel blocker ML418 are in an extended conformation with the cytosolic domain disengaged from the transmembrane domain, as has been observed in the previously obtained unliganded cryo-EM structures of Kir channels. That the channel in the presence of ML418 adopts the extended conformation as observed in apo cryo-EM Kir structures, suggests that the blocked conformation of the channel may be similar to the resting state. We observed the ML418 bound structure with a closed cytosolic exit, G-loop gate, closed helix bundle crossing, and voided selectivity filter. The E276A mutant also adopts the extended conformation and has a partially occupied selectivity filter, closed helix bundle crossing, and closed G-loop gate. As the major difference between the two extended conformation structures is the unoccupied selectivity filter, this could suggest that gating of the selectivity filter occurs at both the extracellular and pore-facing sides of the selectivity filter. In MthK, the selectivity filter does not have to adopt a collapsed conformation to be almost completely impermeable to K^+^ ions [23]. When refined in C1 symmetry the ML418 bound Kir7.1 is observed to adopt differing contacts between subunits within the tetramer. Where contact between Glu149 within the pore and ML418 is observed an altered transmembrane contact network results in the formation of an intersubunit contact between Trp residues that effectively occludes the PIP2 binding site and may represent the desensitized state.

In calcium-gated prokaryotic potassium channel MthK [23], the allosteric communication between the selectivity filter and the activation gate is mediated by contacts between the M2 helix and the selectivity filter S4 site widens the selectivity filter. In both ML418 and E276A structures, the S4 subsite exhibits a network of internal selectivity filter contacts, indicating relative constriction at this site. In the unoccupied state, the selectivity filter exhibits a loss of interactions with the M2 and pore helix that stabilize the ion-occupied state. Both Met125 and Pro127 are conserved at selectivity filter sites +2 and +4 among the voltage-gated channels, but among the Kir channels, they are unique to Kir7.1 [3, 5]. In all other Kir channels, the residue at the equivalent position to Met125 is an arginine. This is also observed for the Kirbac from *Burkholderia pseudomallei*, which also contains the Met and Pro of the voltage-gated channels [16, 17, 27]. Interestingly, near this same position in Kirbac (2WLL), magnesium ions at the extracellular turret loops are observed and coordinated by a glutamine residue. In eukaryotic Kir channels, either glutamine or asparagine are observed at this position, while in Kir7.1 a tyrosine is present. It appears when arginine is present at the +2 position it serves to stabilize interactions between the pore helices and the turret loops and thus may serve to configure an ion binding site in close proximity to the pore entry site. Kir7.1 displays unique rectification properties with no dependence on external potassium, a weak voltage dependence, low sensitivity to extracellular blockage by Ba^2+^ and Cs^2+^, and no dependence on Mg^2+^ for intracellular blockade. Substitution of Met125 with arginine restores Kir7.1 dependence on potassium concentration and sensitivity to divalent cation blockade. Kirbac coordinates magnesium at this site through glutamine side chain carbonyl and additionally by an aspartate residue while in the eukaryotic Kir channels, tyrosine is observed instead of aspartate. Potassium has been shown to prefer pi-cation interactions over electrostatic interactions and perhaps Kir channels have evolved different mechanisms to relay capture of external potassium ions. Interestingly, in the docked conformation of the Kir3.2 cryoEM structure [28], the glutamine residue is oriented in the same manner as in the magnesium-bound Kirbac [27]. However, in the extended conformation, the glutamine is instead oriented downwards towards the pore helix and drawn away from the vestibule due to an unraveling of the terminal turn of the pore helix. It is possible that Kir channels have evolved the use of both electronegative and aromatic residues to sense external potassium and that subtle changes in the configuration of the turret loops determined by the orientations of residues at or near the selectivity filter affect the ability of the channels to capture external ions.

Transition to the open conducting conformation of Kir7.1 can be induced by PIP_2_. Relative to the extended conformation, there is a rotation of cytosolic domains about the pore axis by ∼24° and intimate contacts formed that bridge the cytosolic G-loop and tether helix to M2 transmembrane and slide helices. The distance between the cytosolic and transmembrane domains is decreased, and this is achieved through an increase in the helical character of both the tether and slide helices. This shortening of the slide helix causes engagement of the N-terminal peptide with the cytosolic domain, switching from interaction with the C-D loop in the extended form to a preference for the ß-hairpin at the subunit interface in the docked conformation. This is reminiscent of the latched and unlatched conformations reported for the KirBac structures [27]. In the PIP_2_ bound structure, Met49 from the slide helix is in the cis conformer and interacts with intersubunit Thr278 from the G-loop and Trp45 from the same slide helix. Trp45, in turn, interacts with the inner M2 helix and the tether helix, thus keeping the intersubunit M2 helices in close proximity, maintaining the helical turns in the tether helix, and bringing the G-loop close to the membrane. Maintenance of these contacts positions Val157 away from the pore at the helix bundle crossing or hydrophobic collar. In the extended conformation, the helices are splayed, and Val157 orients its sidechain into the pore, creating a hydrophobic bandwidth of the pore of ∼5Å which increases to ∼7Å in the docked conformation, sufficient to allow passage of a hydrated potassium ion (6.6Å). The PIP_2_ bound and the R162Q mutant adopt the docked conformation, both seem to be in a conductance state. Ions are observed above and below the G-loop gate, possibly reflecting an inability to traverse the gate at this constriction point. Notably, Kir7.1 constricts at the G-loop in the open conducting configuration, creating a narrow opening lined by glutamate carbonyl groups with a width close to the ionic radius of potassium, suggesting that conduction through the G-loop occurs in a manner analogous to the selectivity filter proceeding through loss of the solvation shell. In the E276A structure, the G-loop is unable to adopt the conductive conformation due to the loss of intersubunit contacts with Arg285. In this case it appears ions are unable to traverse the pore at both the helix bundle crossing and the G-loop gate. This same contact is observed in the cryo-EM structure of the Kir2.1 channel [18], with a loss of intersubunit contacts between R312 and E303 and H221 being attributed to a loss of PIP_2_ sensitivity. It appears that closing of the G-loop gate in Kir7.1 proceeds courter-intuitively through widening the pore and increasing the hydrophobic character of the pore lining residues. Another constriction point exists at the cytosolic exit on the intracellular side of the pore. In the ML418 bound structure, this is constricted and lined by Arg203 thus creating a narrow electro-repulsive barrier to ion conduction. In the extended conformation, the N-terminal peptide interacts with residues from the CD loop that constitute the sodium binding site in the GIRK channels [14]. When docked to the transmembrane region, the N-terminal peptide switches to interact with the beta-hairpin at the subunit interface thus influencing the conformation of the pore at the cytosolic exit. In the open conducting conformation, the pore instead becomes lined by the acidic residues Asp232 and Glu233. These residues are equivalent to the potassium ion binding site occupied by D255 in the Kir2.1 strontium-bound cryo-EM structure [18].

Pharmacological studies have suggested that Kir7.1 can be directly gated by interaction with the MC4R in a G-protein-independent manner [9], with the endogenous MC4R agonist α-MSH resulting in channel closure and the endogenous antagonist opening the channel [9]. However, in vivo data for the physiological coupling of MC4R to Kir7.1 has been lacking. For example, the deletion of Kir7.1 from MC4R neurons produced only modest late-onset effects on body weight [29]. The demonstration here that the channel blocker ML418 binds to Kir7.1, confirming previous pharmacological data, led us to test the in vivo consequences of Kir7.1 blockade on feeding behavior and body weight.

ML418 treatment had no effect on the resting membrane potential of MC4R PVH neurons in a slice preparation but did block α-MSH-induced depolarization of the cells. Peripheral treatment of mice with the compound activated a large number of neurons in the PVH, including around 70% of the MC4R neurons, as indicated by induction of c-Fos expression. Peripheral administration or stereotaxic injection of ML418 into the PVH inhibited food intake at magnitudes comparable to the administration of the MC4R agonist setmelanotide and also induced weight loss, and these effects were dependent on the expression of Kir7.1 in MC4R neurons. Thus, in contrast to tissue specific deletion of Kir7.1 from MC4R neurons, pharmacological blockade of Kir7.1 in MC4R neurons does indeed appear to mimic the effects of MC4R agonists on the activation of MC4R neurons and the subsequent inhibition of food intake and weight loss [30]. Furthermore, the functional coupling of MC4R to Kir7.1 does appear to have physiological consequences, as this signaling complex appears to mediate the hyperphagia and weight gain in response to antipsychotic drugs like clozapine and risperidone (Chen Liu, personal communication).

The demonstration here that fusion proteins containing both MC4R and Kir7.1 retain MC4R-mediated regulation of Kir7.1 conductance also strongly argues for the direct coupling of MC4R to Kir7.1. Kir7.1 displays parallels to the voltage-gated channels in the selectivity filter, and it is possible that they may also share some features of the gating mechanism as well. The voltage-gated channels have, in addition to the usual K^+^ channel pore architecture, a four-helix transmembrane domain known as the voltage sensing domain that is connected to the pore helices with a linker helix that lies parallel to the membrane in a manner similar to the slide helix in the Kir channels. Vertical movement of the S4 helix of the voltage sensing domain tugs on the linker helix and results in the widening of the pore helices and opening of the channel. The coupling of movement in one transmembrane domain to channel gating has also been observed in complexes of the K_ATP_ channel with the sulfonylurea receptor [8]. Movement of transmembrane helices in SUR1 in response to ligand binding and alternate engagement and disengagement of the N-terminal peptide of K_ATP_ by SUR1 correspond to transitions between docked and extended conformations of the K_ATP_ channel [8]. Whether similar interactions exist between MC4R and Kir7.1 remains to be addressed.

## METHODS

### Expression and purification

Full-length wild-type and mutant human Kir7.1 constructs were cloned into pFastBac vector encoding an N-terminal GB1 tag and TEV protease cleavage site and expressed in Sf9 insect cells. Cell membranes were solubilized in 50 mM Hepes, pH 7.5, 150 mM KCl, 2% DDM and 0.2% CHS. Supernatant was incubated overnight at 4°C with IgG sepharose beads. Beads were collected and washed with 50 mM Hepes, pH 7.5, 150 mM KCl, 0.1% DDM and 0.1% CHS. Resin was exchanged into MNG detergent on column prior to overnight incubation with TEV protease. Unbound material was collected and passed over a TALON affinity column. Unbound fractions were collected, concentrated and loaded on S200 superdex gel filtration column equilibrated in 50 mM Hepes, pH7.5, 150 mM KCl, 0.005% MNG and 0.0005% CHS. Peak fractions were concentrated to 6-8 mg/mL.

### Sample preparation and data collection

A volume of 3.5 μL of the protein sample was applied to glow-discharged C-flat 1.2/1.3 400 mesh Au grids, blotted for 3.5 s at room temperature and 100% humidity and plunge-frozen in liquid ethane using a Vitrobot Mark IV (FEI). Micrographs were collected on a Titan Krios G4i microscope operated at 300 kV and equipped with K3 direct electron detector. Data collection was automated with EPU and SerialEM. Movies were recorded in electron-counting mode using exposures of 3 seconds with a total dose of ∼60 electrons/Å.

### Data processing and refinement

The movies were motion-corrected and dose-weighted using MotionCor2, and contrast function parameters were estimated using CTFFIND4. Particle picking was first performed using Blob picker implemented in CryoSparc and extracted particles were subjected to one round of 2D classification. Selected 2D classes were then used for particle picking using Template picker in CryoSparc. The extracted particles were then passed through three rounds of 2D classification and selection. Particles from selected classes were then used to generate ab initio models without applied symmetry. Heterogenous refinement of ab initio classes with C4 symmetry was performed, followed by non-uniform refinement in Cryosparc. Initial models for wild-type and mutant Kir7.1 generated using alphafold were placed in cryo-EM maps using Dock in Map implemented in Phenix followed by rounds of real space refinement.

### Transfection and cell culture

HEK293T cells (ATCC, CRL-11268) were maintained in Dulbecco’s modified Eagle’s medium (Invitrogen), supplemented with 10% fetal bovine serum (Life Technologies, Carlsbad, CA, USA), and 10% Antibiotic-Antimycotic (Gibco), and cultured at 37°C in a humidified 5% CO2 incubator. For electrophysiology experiments, 2.5×105 cells were seeded in 100 mm culture dishes and transfected 24 hours later with 4 μg of WT or mutant/MC4R-Kir7.1 tethered constructs cDNA, using X-tremeGENE HP DNA transfection Reagent (Roche Diagnostics, Basel, Switzerland) following the manufacturer’s instructions. Recordings were conducted after 48 hours. Stably transfected T-REx-HEK-293 cell lines expressing Kir7.1-M125R or MC4R-Kir7.1-M125R were utilized for ML418 recording experiments. Kir channel expression was induced by culturing cells overnight in media containing 1 µg/ml doxycycline.

### Cell electrophysiological experiments

Whole-cell patch clamp recordings were performed following the standard procedure outlined by the manufacturer for the SyncroPatch 384PE system (Nanion Technologies, Munich, Germany). Cells, suspended at a concentration of 400,000 cells/ml, were prepared in an external solution containing 140 mM NaCl, 2 mM CaCl2, 4 mM KCl, 1 mM MgCl2, 5 mM glucose, and 10 mM HEPES (pH 7.4 with NaOH, 298 mOsm). These cells were then loaded into a multi-hole (4 holes per well) 384-well Nanion patch clamp (NPC) chip, featuring a thick borosilicate glass bottom with an average resistance pore size of ∼4 mΩ. Electrophysiological protocols were applied using the built-in functionality of the SyncroPatch 384PE system, and data acquisition was performed using the PatchControl 384 application (Nanion Technologies). Parameters such as seal resistance, capacitance, and series resistance were determined for each well following the application of a test pulse and monitored throughout the experiment. Upon achieving whole-cell configuration, the cells were internally filled with a solution containing 10 mM KCl, 10 mM NaCl, 110 mM KF, 10 mM EGTA, and 10 mM HEPES (pH 7.2 with KOH, 284 mOsm). Data analysis was conducted using the DataControl 384 software (Nanion Technologies) and GraphPad Prism (GraphPad Software version 9.4).

### Experimental animal models

*Animals (mice).* The University of Michigan Institutional Animal Care and Use Office approved all animal experiments. Experiments were performed in male C57BL/6J (wild-type, WT) purchased from The Jackson Laboratory (JAX), transgenic *Mc4r-2a-Cre (JAX strain #:* 030759, bred in-house), transgenic Kcnj13^flox/flox^ described and validated in [29] and bred in-house. All experimental mice were 8-14 weeks old except for the dietary-induced obese (DIO) mice, which were 30-32 weeks old. All mice were single-housed with a 12-h light/12-h dark cycle and fed either regular chow (5LOD from Lab Diet) or high-fat diet (60 kcal% fat, Research Diets, Cat#: D12492). Mice had free access to food and water unless specified. DIO models were developed by exposing mice to a high-fat diet at 4–5 weeks of age for 16–20 weeks. Data from individual mice are shown in the figures.

### Drugs and reagents

ML418 was first synthesized and characterized in [12]. A-4166 and VU0134992 (Tocris) were dissolved in sterile DMSO and subsequently in USP-grade sterile saline. Setmelanotide (Rhythm Pharmaceuticals) was dissolved in sterile saline, and working concentrations were made on the day of the experiment using Ringer’s solution in mM: 123 NaCl, 1.5 CaCl_2_ 5 KCl, pH 7.3-7.4.

### Slice electrophysiology

Acute hypothalamic brain slices were prepared similarly to previously described [9]. Briefly, adult male and female MC4R-GFP [31] mice were anesthetized with isoflurane and decapitated. The brain was removed and submerged in carbogen (95% O_2_ and 5% CO_2_) saturated with ice-cold sucrose solution containing (in mM) 126.2 NaCl, 3.1 KCl, 2 CaCl2, 1 MgCl2, 1 NaH2PO4, 26.2 NaHCO3, 10 glucose and 11 sucrose (320 mosm/kg, pH 7.39). Brain blocks containing the hypothalamus were made, and slices of 200 μm thickness were cut. Slices were transferred to a glass beaker containing oxygenated ACSF at 31 °C for 1 hour. Slices were placed in a recording chamber with warmed (31-32 °C) ACSF at a flow rate of 2-3 ml/min. GFP-PVN cells were patched using epifluorescence and IR-DIC optics. This study used whole-cell current clamp recordings to obtain membrane potential on PVN MC4R neurons. Recordings were performed using patch pipettes of 3 - 5 MΩ resistance when filled with a solution containing (in mM) 125 K gluconate, 8 KCl, 4 MgCl2, 10 HEPES, 5 NaOH, 4 Na_2_ATP, 0.4 Na_3_GTP, 5 Na_2_-creatine phosphate, 7 sucrose and 7 KOH which resulted in a pH ∼7.23 and osmolality of 295–300 mosmol/kg. α-MSH (250 nM) was bath applied to the slice. To examine the effect of ML418 on α-MSH-induced effects on membrane potential, slices were pre-incubated with ML418 (5µm). Data were acquired at 10 kHz using a MultiClamp 700A amplifier (2000X gain, −3dB filter freq: 5 kHz) and Clampex 10.0.1 software (Axon Instruments, Union City, CA).

### Acute feeding experiments

Mice were individually housed for at least 7 days and acclimated to handling and saline injection for 5-7 days. Compound/drug injections occurred at least 30 minutes prior to the onset of the dark cycle. ML418 (5-7.5 mg/kg) and A-4166 (5 mg/kg) were intraperitoneally (ip) injected into mice at the onset of the dark cycle. ML418 and VU0134992 (500pmoles/1uL) were delivered into the left lateral ventricle (icv). Setmelanotide and ML418 (200pmoles/0.5uL/side) were injected into the paraventricular nucleus of the hypothalamus (PVH) of mice. Pre-weighed food pellets were given to mice for all experiments, and food intake was measured over 24 hours or at the 24-hour mark after compound administration. Body weight and water intake were measured 24 hours post-administration.

### Surgery and cannula placement

Mice were housed in groups of 3 or more with ad libitum access to standard chow and water before surgery. An Intracerebroventricular (icv) cannula was implanted under 3-4% (v/v) isoflurane before being placed in a stereotaxic surgical frame (David Kopf Instruments, Tujuna, CA) and then maintained at 1.5-2% (v/v) isoflurane for the rest of the surgery. Cannula were implanted as previously described [32]. The skull was exposed by an incision and leveled between lambda and bregma. Guide Cannula (Plastics One, Roanoke, VA) targeted the left lateral ventricle using the following flat skull coordinates (-0.460 mm posterior to Bregma, 1.00 lateral to the midline, and -2.20 ventral to the surface of the skull). The cannula was secured to the skull with dental cement. Following recovery, mice received an icv injection of 20 ng angiotensin II (Sigma-Aldridge) to confirm cannula placement by angiotensin-induced water intake. Cannula placement was also confirmed postmortem under a microscope. For intra-PVN cannulation, bilateral guide cannula (Plastics One, Roanoke, VA) dipped in DiD dye Invitrogen) targeted the PVN at the coordinates (0.7 mm posterior to bregma, ±0.3 mm from midline, 5.0 mm ventral to the surface of the skull). The proper coordinates were histologically verified post-mortem by visualizing DiD under a fluorescent microscope. Only mice with DiD expression in the PVH were included in the final analysis.

### Immunofluorescence staining

MC4R-GFP mice were handled and peripherally injected with 150 μl sterile daily for 4-5 days before the experiment to minimize background neuronal activation (C-fos) due to stress. Mice received an injection of 5mg/kg ML418 (ip) or vehicle (DMSO, 20uL, ip). After 90 mins, mice were given a lethal dose of anesthesia and transcardially perfused with 4% paraformaldehyde in 0.1 M phosphate buffer (pH 7.4). Brains were removed and postfixed in the same fixative overnight. Tissues were rinsed in PBS and cryoprotected in 30% sucrose in PBS until the tissue sank. Brains were embedded in OCT (Tissue-Tek), frozen in a dry ice ethanol bath, and stored at -80 °C. Hypothalamic sections were cut on a cryostat at 50 μm thick. The tissues were immediately washed with PBS, rocking slowly at room temperature (RT) to warm. Free-floating sections were blocked for 1 h with 0.3% triton and 5% normal serum in PBS. Primary cFos (1:2000; synaptic systems 226 004) and primary GFP antibody (1:4000; Aves Labs GFP-1020) were diluted in the blocking buffer. Sections were incubated with primary antibodies overnight, rocking at RT, rinsed 3 times in PBS, and incubated with secondary antibodies in blocking buffer (Alexa conjugated antibodies, Thermo Fisher Scientific, 1:500) for 2 h at RT. The primary antibodies were omitted for control experiments, and sections were incubated with the blocking buffer instead. Sections were rinsed 3 times with PBS, counterstained with Hoechst 33342 (Thermo Fisher Scientific, 1/2000 in PBS-0.05% Tween 20), and rinsed 2 times with PBS. Sections were moved and slides and coverslips were mounted with Fluoromount G (Southern Biotechnology). C-Fos was counted on sections on all PVN sections (0.60 -1.20 mm posterior to bregma).

## ACKNOWLEDGMENTS

RDC was supported by NIH grants RO1 DK110403 and DK070332, and NSD was supported by NIH T32 DK101357. Electrophysiological studies were conducted in part using a Nanion Synchropatch device supported by NIH S10OD025203 (RDC). We thank members of the Life Sciences Institute Center for Chemical Genomics and Cryoelectron Microscopy Core.

## AUTHOR CONTRIBUTIONS

Author contributions: AP, CCH, NSD, MG-L, LEG, and RDC designed the experiments. AP, CCH, NSD, MG-L, LK, AR, MS, XJ and LEG collected data. AP, CCH, NSD, MG-L, MS, XJ, LEG, and

RDC analyzed the data, and AP, CCH, NSD, LEG, and RDC wrote the manuscript.

## DECLARATION OF INTERESTS

RDC, LEG, and the University of Michigan have equity in Courage Therapeutics, and RDC serves on the company board. The other authors declare no competing interests.

## REFERENCES

1. Hibino H, Inanobe A, Furutani K, Murakami S, Findlay I, Kurachi Y. Inwardly rectifying potassium channels: their structure, function, and physiological roles. Physiol Rev. 2010;90(1):291–366. doi: 10.1152/physrev.00021.2009.

2. González C, Baez-Nieto D, Valencia I, Oyarzún I, Rojas P, Naranjo D, Latorre R. K(+) channels: function-structural overview. Compr Physiol. 2012;2(3):2087–149. doi: 10.1002/cphy.c110047.

3. Döring F, Derst C, Wischmeyer E, Karschin C, Schneggenburger R, Daut J, Karschin A. The epithelial inward rectifier channel Kir7.1 displays unusual K+ permeation properties. J Neurosci. 1998;18(21):8625–36. doi: 10.1523/jneurosci.18-21-08625.1998.

4. Shimura M, Yuan Y, Chang JT, Zhang S, Campochiaro PA, Zack DJ, Hughes BA. Expression and permeation properties of the K(+) channel Kir7.1 in the retinal pigment epithelium. J Physiol. 2001;531(Pt 2):329–46. doi: 10.1111/j.1469-7793.2001.0329i.x.

5. Krapivinsky G, Medina I, Eng L, Krapivinsky L, Yang Y, Clapham DE. A novel inward rectifier K+ channel with unique pore properties. Neuron. 1998;20(5):995–1005. doi: 10.1016/s0896-6273(00)80480-8.

6. Hejtmancik JF, Jiao X, Li A, Sergeev YV, Ding X, Sharma AK, et al. Mutations in KCNJ13 cause autosomal-dominant snowflake vitreoretinal degeneration. Am J Hum Genet. 2008;82(1):174–80. doi: 10.1016/j.ajhg.2007.08.002.

7. Sergouniotis PI, Davidson AE, Mackay DS, Li Z, Yang X, Plagnol V, et al. Recessive mutations in KCNJ13, encoding an inwardly rectifying potassium channel subunit, cause leber congenital amaurosis. Am J Hum Genet. 2011;89(1):183–90. doi: 10.1016/j.ajhg.2011.06.002.

8. Martin GM, Yoshioka C, Rex EA, Fay JF, Xie Q, Whorton MR, et al. Cryo-EM structure of the ATP-sensitive potassium channel illuminates mechanisms of assembly and gating. Elife. 2017;6. doi: 10.7554/eLife.24149.

9. Ghamari-Langroudi M, Digby GJ, Sebag JA, Millhauser GL, Palomino R, Matthews R, et al. G-protein-independent coupling of MC4R to Kir7.1 in hypothalamic neurons. Nature. 2015;520(7545):94-8. doi: 10.1038/nature14051.

10. Jiang Y, Lee A, Chen J, Cadene M, Chait BT, MacKinnon R. The open pore conformation of potassium channels. Nature. 2002;417(6888):523-6. doi: 10.1038/417523a.

11. Jiang Y, Lee A, Chen J, Cadene M, Chait BT, MacKinnon R. Crystal structure and mechanism of a calcium-gated potassium channel. Nature. 2002;417(6888):515-22. doi: 10.1038/417515a.

12. Swale DR, Kurata H, Kharade SV, Sheehan J, Raphemot R, Voigtritter KR, et al. ML418: The First Selective, Sub-Micromolar Pore Blocker of Kir7.1 Potassium Channels. ACS Chem Neurosci. 2016;7(7):1013-23. doi: 10.1021/acschemneuro.6b00111.

13. Hansen SB, Tao X, MacKinnon R. Structural basis of PIP2 activation of the classical inward rectifier K+ channel Kir2.2. Nature. 2011;477(7365):495-8. doi: 10.1038/nature10370.

14. Whorton MR, MacKinnon R. Crystal structure of the mammalian GIRK2 K+ channel and gating regulation by G proteins, PIP2, and sodium. Cell. 2011;147(1):199-208. doi: 10.1016/j.cell.2011.07.046.

15. Rohács T, Lopes CM, Jin T, Ramdya PP, Molnár Z, Logothetis DE. Specificity of activation by phosphoinositides determines lipid regulation of Kir channels. Proc Natl Acad Sci U S A. 2003;100(2):745–50. doi: 10.1073/pnas.0236364100.

16. Kuo A, Gulbis JM, Antcliff JF, Rahman T, Lowe ED, Zimmer J, et al. Crystal structure of the potassium channel KirBac1.1 in the closed state. Science. 2003;300(5627):1922-6. doi: 10.1126/science.1085028.

17. Kuo A, Domene C, Johnson LN, Doyle DA, Vénien-Bryan C. Two different conformational states of the KirBac3.1 potassium channel revealed by electron crystallography. Structure. 2005;13(10):1463–72. doi: 10.1016/j.str.2005.07.011.

18. Fernandes CAH, Zuniga D, Fagnen C, Kugler V, Scala R, Péhau-Arnaudet G, et al. Cryo-electron microscopy unveils unique structural features of the human Kir2.1 channel. Sci Adv. 2022;8(38):eabq8489. doi: 10.1126/sciadv.abq8489.

19. Shinkai H, Nishikawa M, Sato Y, Toi K, Kumashiro I, Seto Y, et al. N-(cyclohexylcarbonyl)-D-phenylalanines and related compounds. A new class of oral hypoglycemic agents. 2. J Med Chem. 1989;32(7):1436-41. doi: 10.1021/jm00127a006.

20. Villanueva S, Burgos J, Lopez-Cayuqueo KI, Lai KM, Valenzuela DM, Cid LP, Sepulveda FV. Cleft Palate, Moderate Lung Developmental Retardation and Early Postnatal Lethality in Mice Deficient in the Kir7.1 Inwardly Rectifying K+ Channel. PLoS One. 2015;10(9):e0139284. doi: 10.1371/journal.pone.0139284.

21. Kharade SV, Kurata H, Bender AM, Blobaum AL, Figueroa EE, Duran A, et al. Discovery, Characterization, and Effects on Renal Fluid and Electrolyte Excretion of the Kir4.1 Potassium Channel Pore Blocker, VU0134992. Mol Pharmacol. 2018;94(2):926-37. doi: 10.1124/mol.118.112359.

22. Markham A. Setmelanotide: First Approval. Drugs. 2021;81(3):397-403. doi: 10.1007/s40265-021-01470-9.

23. Kopec W, Rothberg BS, de Groot BL. Molecular mechanism of a potassium channel gating through activation gate-selectivity filter coupling. Nat Commun. 2019;10(1):5366. doi: 10.1038/s41467-019-13227-w.

24. Klement G, Nilsson J, Arhem P, Elinder F. A tyrosine substitution in the cavity wall of a k channel induces an inverted inactivation. Biophys J. 2008;94(8):3014–22. doi: 10.1529/biophysj.107.119842.

25. Wang S, Bondarenko VE, Qu Y, Morales MJ, Rasmusson RL, Strauss HC. Activation properties of Kv4.3 channels: time, voltage and [K+]o dependence. J Physiol. 2004;557(Pt 3):705–17. doi: 10.1113/jphysiol.2003.058578.

26. Cuello LG, Jogini V, Cortes DM, Pan AC, Gagnon DG, Dalmas O, et al. Structural basis for the coupling between activation and inactivation gates in K(+) channels. Nature. 2010;466(7303):272-5. doi: 10.1038/nature09136.

27. Clarke OB, Caputo AT, Hill AP, Vandenberg JI, Smith BJ, Gulbis JM. Domain reorientation and rotation of an intracellular assembly regulate conduction in Kir potassium channels. Cell. 2010;141(6):1018–29. doi: 10.1016/j.cell.2010.05.003.

28. Niu Y, Tao X, Touhara KK, MacKinnon R. Cryo-EM analysis of PIP(2) regulation in mammalian GIRK channels. Elife. 2020;9. doi: 10.7554/eLife.60552.

29. Anderson EJP, Ghamari-Langroudi M, Cakir I, Litt MJ, Chen V, Reggiardo RE, et al. Late onset obesity in mice with targeted deletion of potassium inward rectifier Kir7.1 from cells expressing the melanocortin-4 receptor. J Neuroendocrinol. 2019;31(1):e12670. doi: 10.1111/jne.12670.

30. Krashes MJ, Lowell BB, Garfield AS. Melanocortin-4 receptor-regulated energy homeostasis. Nat Neurosci. 2016;19(2):206–19. doi: 10.1038/nn.4202.

31. Liu H, Kishi T, Roseberry AG, Cai X, Lee CE, Montez JM, et al. Transgenic mice expressing green fluorescent protein under the control of the melanocortin-4 receptor promoter. J Neurosci. 2003;23(18):7143–54. doi: 10.1523/JNEUROSCI.23-18-07143.2003.

32. Dahir NS, Gui Y, Wu Y, Sweeney PR, Williams SY, Gimenez LE, et al. Inhibition of the melanocortin-3 receptor (MC3R) causes generalized sensitization to anorectic agents. bioRxiv. 2023. doi: 10.1101/2023.12.05.570114.

