## Supplemental Data for "Structure of the Ion Channel Kir7.1 and Implications for its Function in Normal and Pathophysiologic States"

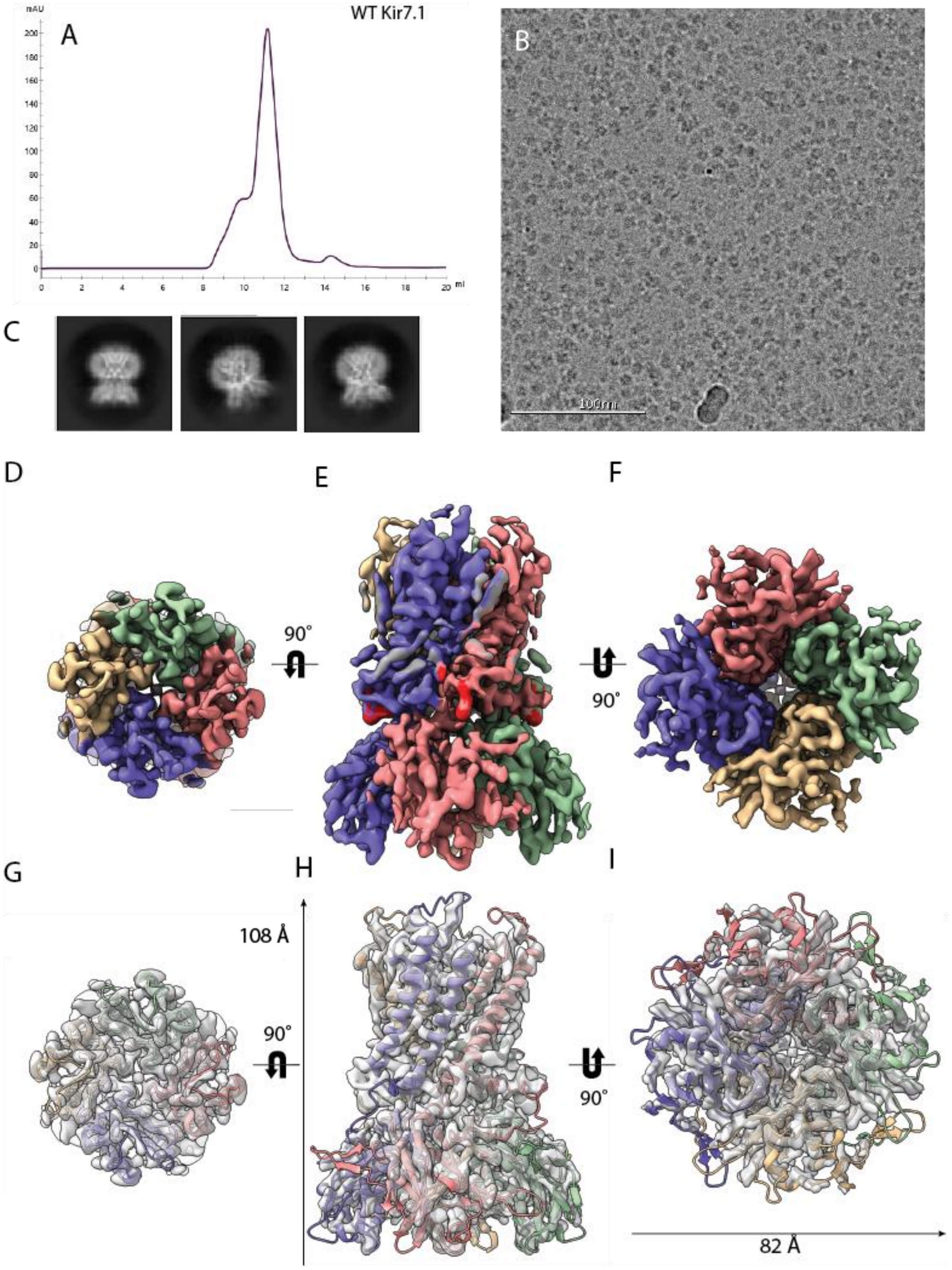


**Figure S1.** A. Superose 6 size exclusion profile of wt Kir7.1 tetramer. B. Cryo-electron micrograph of wt Kir7.1. C. 2D cryo class averages of wt Kir7.1. D-F Cryo-EM volume of wt Kir7.1 bound to PIP2 with surface colored according to chain of fitted tetrameric model. Views down the pore from transmembrane side, side view and from the cytosolic side through the pore are shown. G-I. Cryo-EM volume of wt Kir7.1 bound to PIP2 shown as transparent surface and fitted with tetrameric model colored by individual chains. Views from transmembrane, across membrane and from cytosolic side are shown.

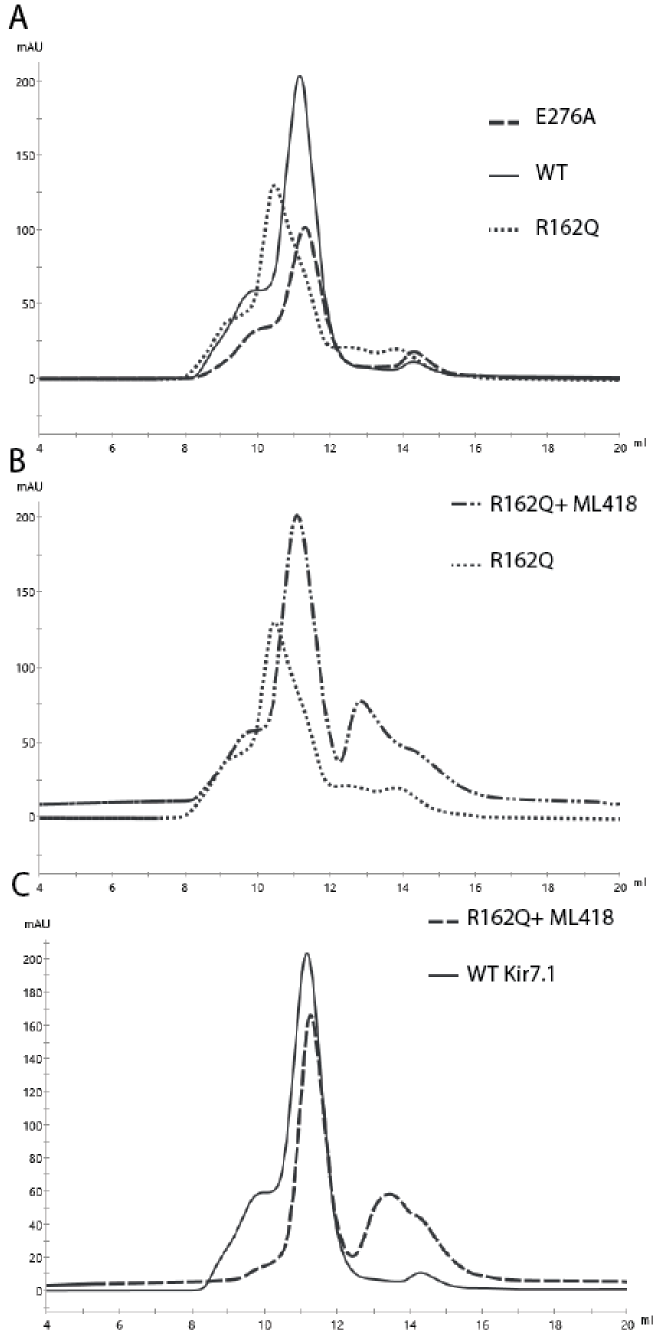


**Figure S2.** A. Overlay of superose 6 size exclusion profiles of apo WT, E276A and R162Q mutant are shown. R162Q displays left shifted profile relative to WT and E276A mutant. B. Overlay of R162Q and R162Q bound with ML418 superose 6 size exclusion profiles. Presence of ML418 shifts R162Q migration to right. C. Overlay of R162Q bound to ML418 and WT Kir7.1 superose 6 size exclusion profile showing equivalent migration position.

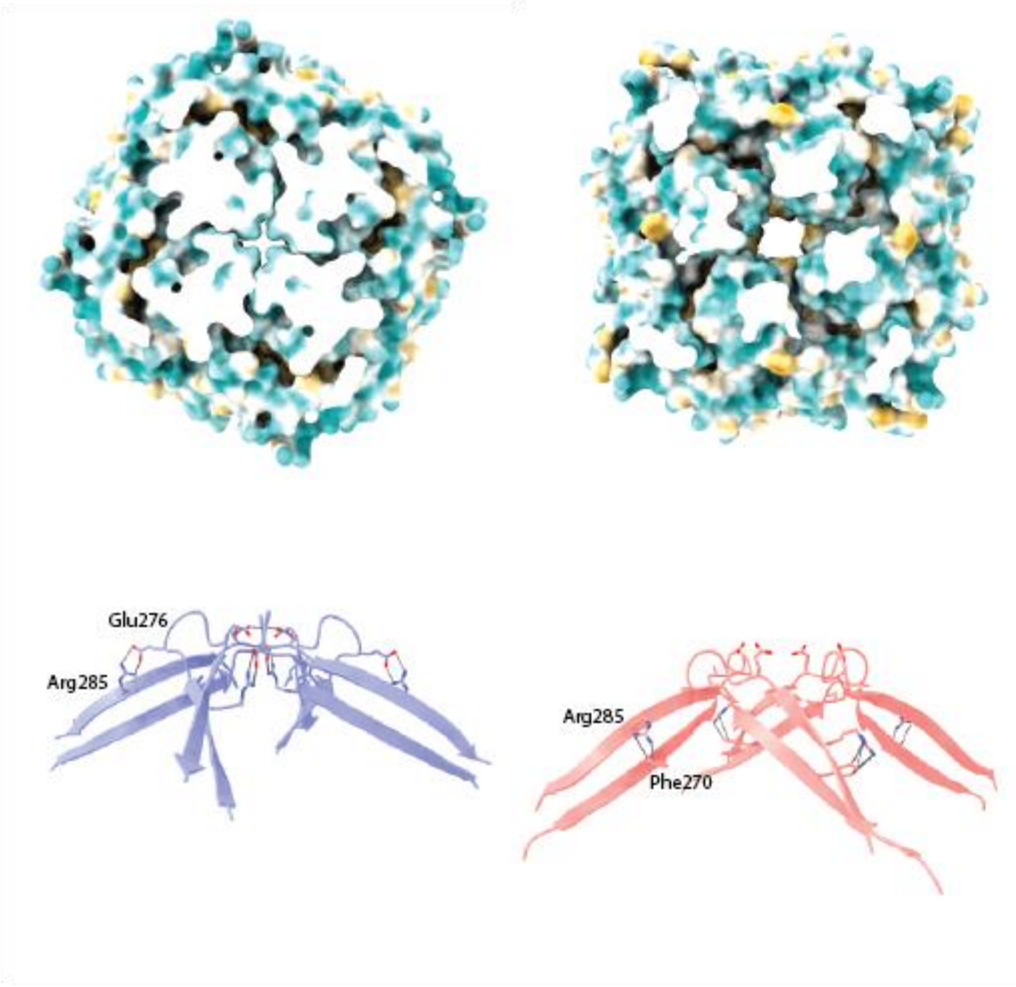


**Figure S3.** Upper panel: View of cytosolic domain looking down the pore from the transmembrane surface above G-loop and rendered as a hydrophobic surface. Width and hydrophobicity are increased in E276A mutant (right) relative to the PIP2 bound WT (left). Lower panel: Side view of G-loop indicating altered interaction of Arg285 in WT (purple) and E276A mutant (salmon). E276A shows loss of stabilization of interdomain contacts in G-loop structure.

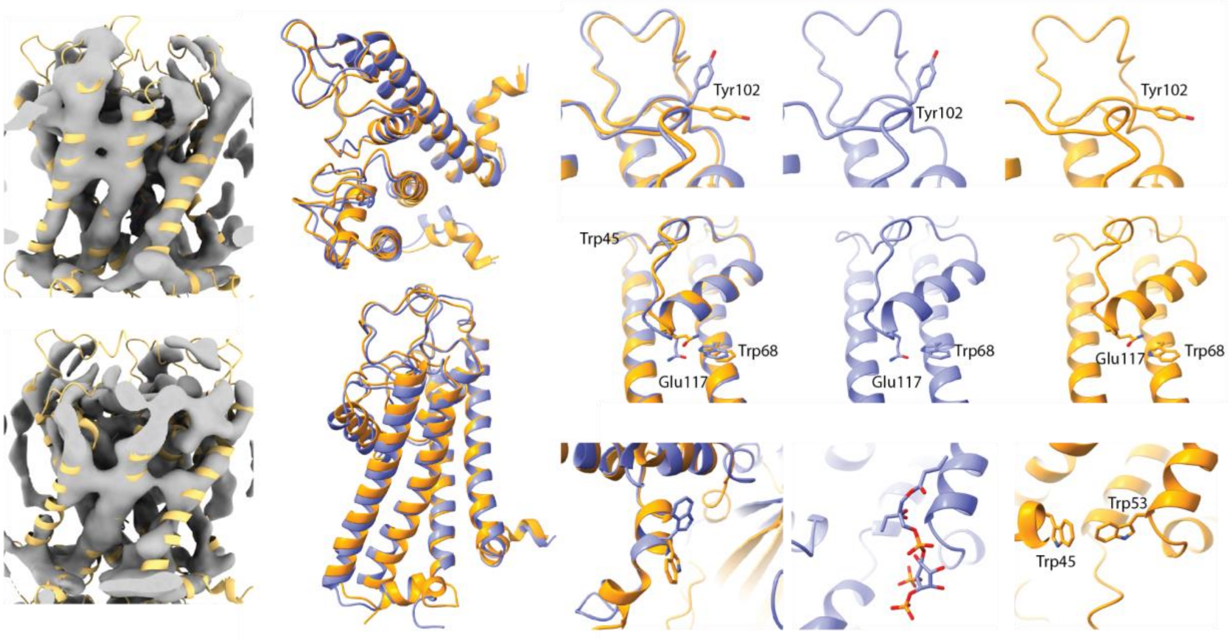


**Figure S4.** Left: Views of volume of C1 symmetry refined density map of ML418 bound Kir7.1 from opposing sides of tetramer. Upper left: continuous density observed between slide helices and contact between M1 helix and pore helix (putative desensitized conformer). Lower left: altered conformation with density corresponding to intersubunit contacts between M1 and M2 helices and a loss of density between slide helices (resting conformer). Upper middle: Alignment of transmembrane domains of ML418 bound Kir7.1 (yellow) and PIP2 bound Kir7.1 (purple). Right panels: Altered residue positions in open conformation vs putative desensitized state: Left aligned, right PIP2 and right ML418; upper flipped Tyr102 in turret region near pore helix, middle: contact gained between Glu117 in pore helix and Trp68 in M1 transmembrane helix, lower: flipped tryptophan in slide helix allows for interaction with adjacent subunit base of M1 helix.

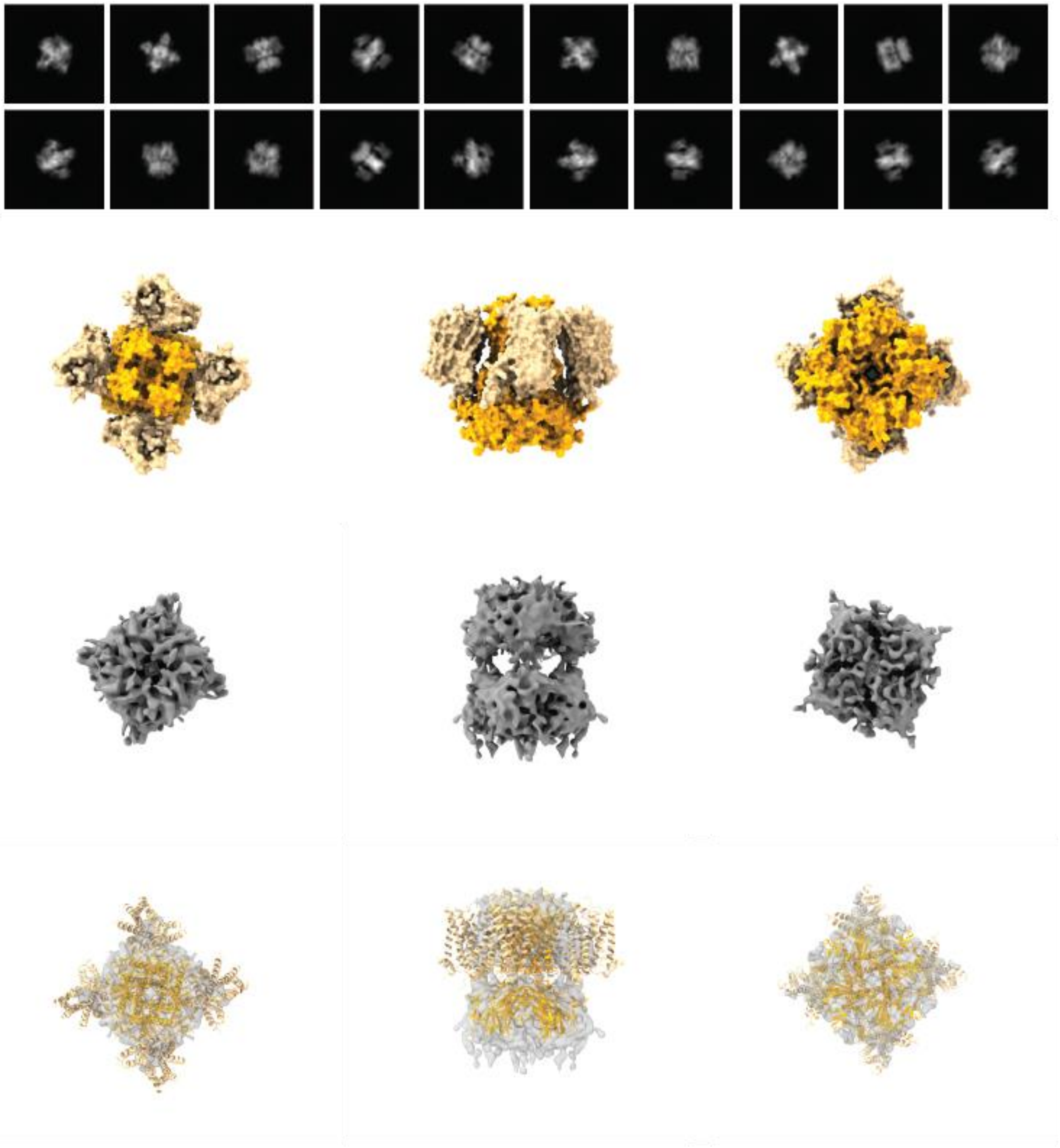

Figure S5. Upper panel - Class averages of simulated particles generated from theoretical volumes from model of MC4R bound to Kir7.1 in extended conformation. ML418 bound Kir7.1 was docked to EM map of linker construct. SHU911 bound MC4R was aligned to alphafold prediction of MC4R:Kir7.1 transmembrane interaction surface. Then the alpha-MSH bound MC4R-G-protein structure (7PIU) was aligned by aligning bound peptides in docking site. This recapitulated tilt angle exhibited by additional density seen attached to Kir7.1 channel. Model is shown as surface, and fitted to experimental volume.

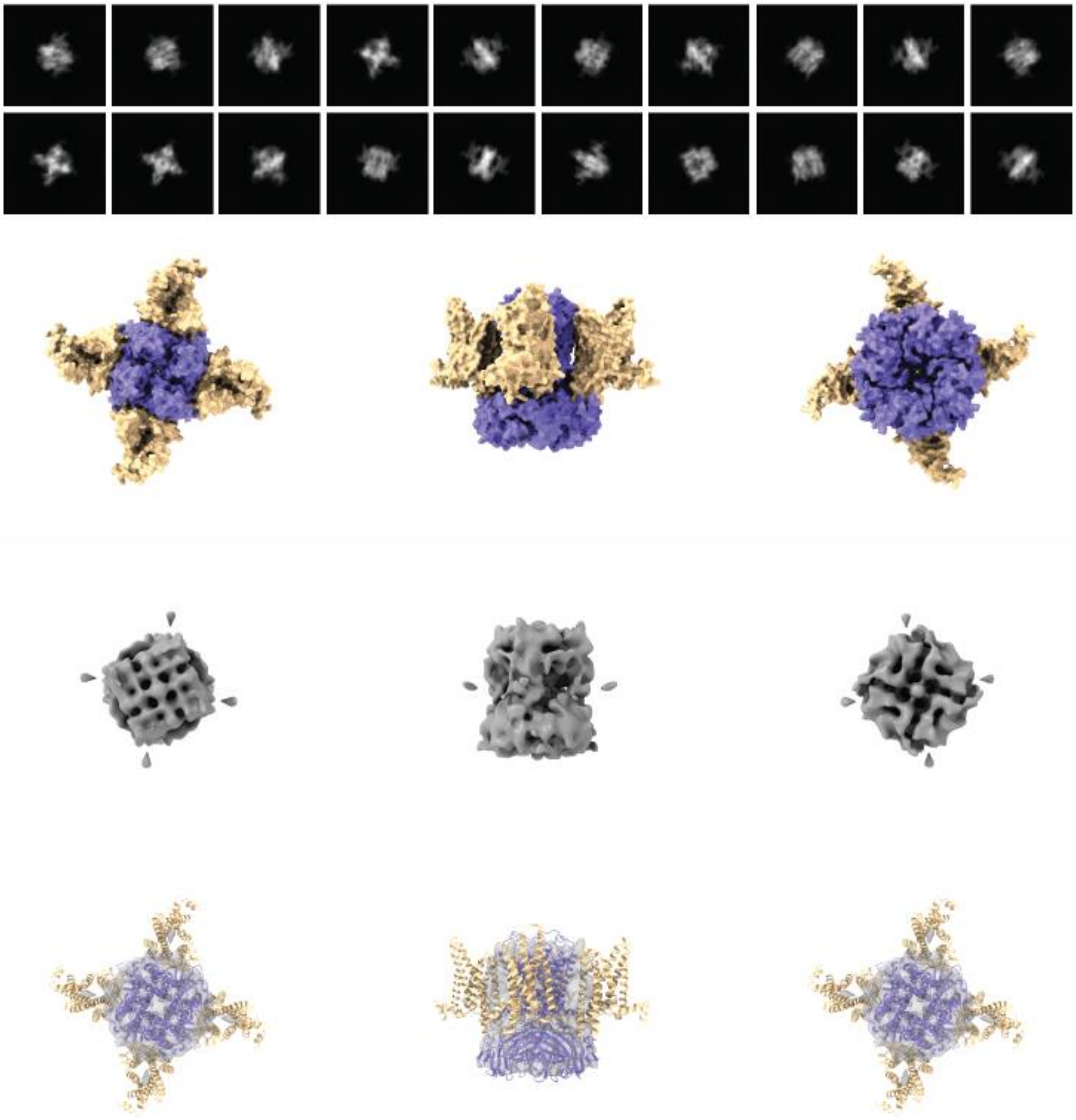

Figure S6. Class averages of simulated particles generated from theoretical volumes from model of MC4R bound to Kir7.1 generated by alphafold. Alphafold model is shown as a surface and fitted to experimental volume.

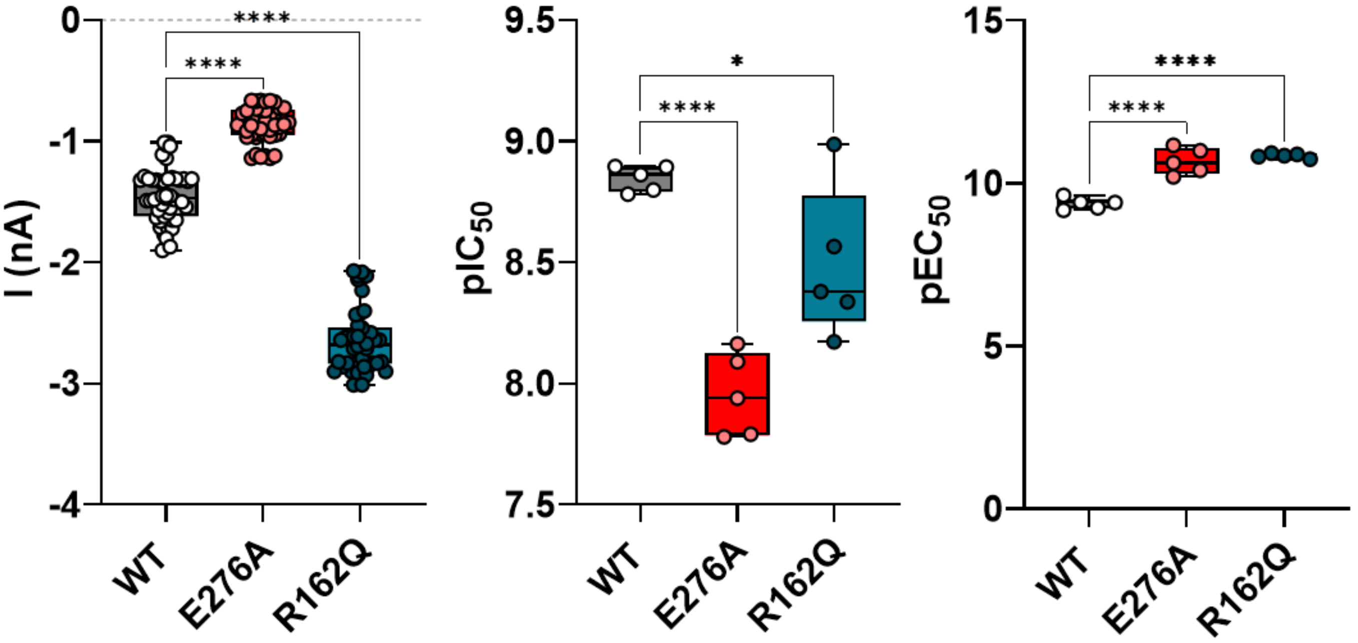


| | Current amplitude (nA ± SEM) | *P* (N) | α-MSH $pIC_{50}$ (± SEM) | *P* (N) | AgRP $pEC_{50}$ (± SEM) | *P* (N) |
|---|---|---|---|---|---|---|
| **WT** | -1.447 ± 0.03246 | (5) | 8.847 ± 0.02387 | (5) | 9.375 ± 0.07919 | (5) |
| **E276A** | -0.8627 ± 0.02236 | <0.0001, (5) | 7.953 ± 0.07734 | <0.0001, (5) | 10.68 ± 0.1808 | <0.0001, (5) |
| **R162Q** | -2.647 ± 0.04071 | <0.0001, (5) | 8.489 ± 0.1394 | 0.0338, (5) | 10.85 ± 0.03234 | <0.0001, (5) |

One-way ANOVA: Dunnett's multiple comparisons test

Figure S7. Bar plots of the effects of Kir7.1 mutants on current amplitude and MC4R-Kir7.1 coupling. The table summarizes the results shown on the bar plots.

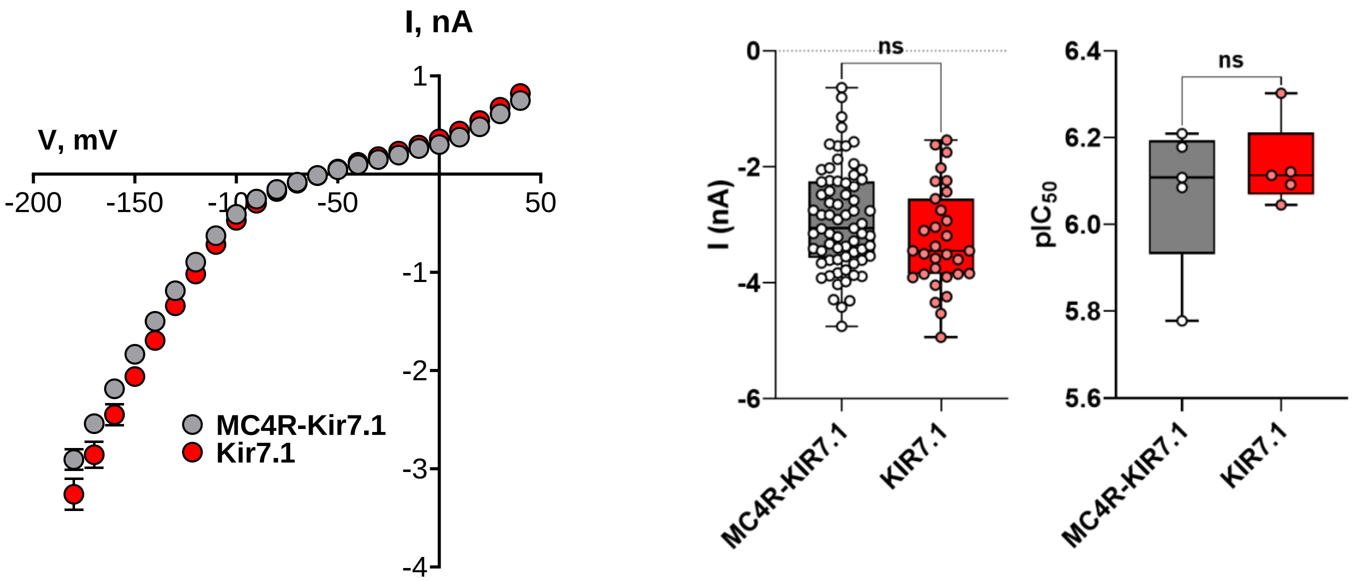


| | Current amplitude (nA ± SEM) | *P* (N) | ML418 pIC$_{50}$ (± SEM) | *P* (N) |
|---|---|---|---|---|
| **MC4R-KIR7.1** | -2.907 ± 0.1068 | (69) | 6.072 ± 0.0767 | (5) |
| **KIR7.1** | -3.2607 ± 0.1568 | 0.0674 (31) | 6.135 ± 0.0438 | 0.4965 (5) |

Unpaired t-test

Figure S8. Top left, Current-voltage plots for Kir7.1 expressed alone or coexpressed with MC4R. Peak currents for each voltage were measured from a holding potential of −60 mV, and step pulses from −180 to 40 mV were applied to the cells in 10 mV increments for 200 ms. Top right, Bar plots of the effects of Kir7.1 expressed alone or coexpressed with MC4R on current amplitude and MC4R-Kir7.1 coupling. Bottom, the table summarizes the results shown on bar plots.

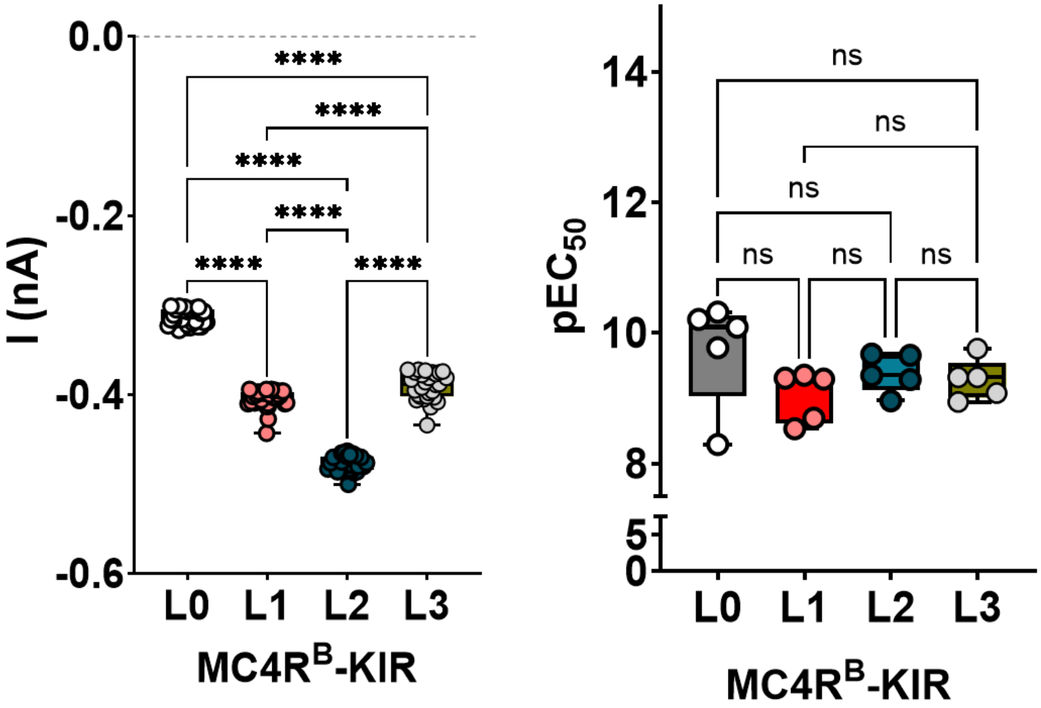


| | | Current amplitude (pA ± SEM), N=28 | One-way ANOVA: Tukey's multiple comparisons test | | | | AgRP pEC$_{50}$ (± SEM), N=5 | One-way ANOVA: Tukey's multiple comparisons test | | | |
|---|---|---|---|---|---|---|---|---|---|---|---|
| | | | | ***P*** | | ***P*** | | | ***P*** | | ***P*** |
| **MC4R$^B$-KIR** | **L0** | -0.3134 ± 0.001661 | L0 vs. L1 | <0.0001 | L1 vs. L3 | <0.0001 | 9.737 ± 0.3727 | L0 vs. L1 | 0.1708 | L1 vs. L3 | 0.8651 |
| | **L1** | -0.4049 ± 0.002015 | L0 vs. L2 | <0.0001 | L2 vs. L3 | <0.0001 | 9.037 ± 0.1748 | L0 vs. L2 | 0.6999 | L2 vs. L3 | 0.9886 |
| | **L2** | -0.4776 ± 0.001691 | L0 vs. L3 | <0.0001 | | | 9.387 ± 0.1319 | L0 vs. L3 | 0.5129 | | |
| | **L3** | -0.3905 ± 0.002912 | L1 vs. L2 | <0.0001 | | | 9.285 ± 0.1389 | L1 vs. L2 | 0.7003 | | |

Figure S9. Bar plots of the effects of the MC4R-Kir7.1 tethered constructs on current amplitude and MC4R-Kir7.1 coupling. The table summarizes the results shown on bar plots.
